# Sialylation of Asparagine 612 inhibits Aconitase activity during mouse sperm capacitation; A possible mechanism for the switch from oxidative phosphorylation to glycolysis

**DOI:** 10.1101/2020.04.28.067298

**Authors:** Ana Izabel Silva Balbin Villaverde, Rachel Ogle, Peter Lewis, Louise Hetherington, Vince Carbone, Tony Velkov, Jacob Netherton, Mark A. Baker

## Abstract

After ejaculation, mammalian spermatozoa must undergo a process known as capacitation in order to successfully fertilize the oocyte. Several post-translational modifications occur during capacitation, including sialylation, which despite being limited to a few proteins, seems to be essential for proper sperm-oocyte interaction. Regardless of its importance, to date, no single study has ever identified nor quantified which glycoproteins bearing terminal sialic acid (Sia) are altered during capacitation. Here we characterize sialylation during mouse sperm capacitation. Using tandem mass spectrometry coupled with liquid chromatography (LC-MS/MS), we found 142 non-reductant peptides, with 9 of them showing potential modifications on their sialylated oligosaccharides during capacitation. As such, N-linked sialoglycopeptides from C4b-binding protein, endothelial lipase (EL), serine proteases 39 and 52, testis-expressed protein 101 and zonadhesin were reduced following capacitation. In contrast, mitochondrial aconitate hydratase (aconitase; ACO2) was the only protein to show an increase in Sia content during capacitation. Interestingly, while the loss of Sia within EL (N62) was accompanied by a reduction in its phospholipase A_1_ activity, the increase of sialylation in the ACO2 (N612) also resulted in a decrease of the activity of this TCA cycle enzyme. The latter was confirmed by N612D recombinant protein with both His and GFP tag, in which the N612D mutant had no activity compared to WT when protein. Computer modelling show that N612 sits atop the catalytic site of ACO2. The introduction of sialic acid causes a large confirmation change in the alpha helix, essentially, distorting the active site, leading to complete loss of function. These findings suggest that the switch from oxidative phosphorylation, over to glycolysis that occurs during capacitation may come about through sialylation of ACO2.

## Introduction

Sperm capacitation is a phenomenon first described by Chang, who demonstrated that freshly ejaculated spermatozoa are unable to fertilize the oocyte immediately [1]. Rather, a period within the female reproductive tract was required [1]. During this period, it is evident that spermatozoa undergo a series of biochemical and metabolic changes to become fully capable of fertilizing the egg [2–4]. From a metabolic perspective, Fraser and Lane first described the phenomenon of a metabolic switch that occurs during capacitation [5]. In this context, freshly ejaculated spermatozoa have high rates of oxygen consumption, however, during *in vitro* capacitation, mouse spermatozoa decrease their reliance on oxidative phosphorylation and switch over to a glycolytic pathway [5]. Using different metabolic substrates, this “switch” was shown to be necessary to achieve fertilization [5]. In addition, the reliance on glycolysis explain why sperm-specific glycolytic knockout mice, including GADPH [6], are infertile due to poor motility. From a biochemical perspective, it is unknown how sperm make this switch. However, considering that spermatozoa are transcriptionally and translationally silent, with the exception of mitochondrial proteins [7], we reasoned that post-translational modifications of existing proteins would likely play a role [8].

Whilst the role of phosphorylation in sperm capacitation is well studied [9–11], few studies have looked at the specific role of protein glycosylation. The latter is characterized by the addition of oligosaccharide side chains via a covalent linkage either at the asparagine (Asn) residue (N-linked) or at the serine/threonine (Ser/Thr) residues (O-linked). There are a number of variations in terms of glycan structures, which is translated in a wide range of biological functions [12, 13]. Notably, terminal sugar sequences are among the features suggested as mediators of more specific roles [12]. In this context, sialic acid (Sia), a monosaccharide with a nine-carbon backbone and negatively charged, is considered an important terminal sugar.

The presence of Sia residues in glycoconjugates at the sperm surface seems to be vital for the success of fertilization. Indeed, in both humans [14] and boars [15] with unexplained infertility or subfertility, a decreased affinity for the lectin wheat germ agglutinin (WGA) has been reported, suggesting either N-acetyl glucosamine and Sia residues are lacking within these cells. Interestingly, surface Sia residues have also been shown to help sperm to survive and migrate inside the female reproductive tract by reducing their phagocytosis and antigenicity [16–18] and by helping them to penetrate the cervical mucus [19]. During sperm capacitation, it has been suggested that Sia residues are shed from the surface of spermatozoa. In this context, studies using lectin binding [20–23], surface charge determination [24, 25] radiolabeling of terminal sialyl residues [26] and HPLC measurement following acid hydrolysis [27] have all indicated that surface sialoglycoconjugates are probably lost or modified when spermatozoa are incubated under capacitating conditions. One model that has been put forward is that the shedding of Sia residues may be triggered by the release of neuraminidases present at the sperm surface [27]. If this is the case, then the presence of oviduct fluid components [21] heparin [22] and/or albumin [28] in the medium probably facilitate this release [27].

Despite the data that show changes in sperm sialylation content during capacitation, no single publication has ever look at which proteins are sialylated in spermatozoa, yet alone quantified their levels during capacitation. With this in mind, we used titanium dioxide (TiO2) followed by peptide-N-glycosidase F (PNGase F) cleavage to search for N-linked Sia-containing peptides and to quantify them during *in vitro* capacitation of mouse sperm. In addition, we further studied both mitochondrial aconitate hydratase (aconitase; ACO2) and endothelial lipase (EL) proteins to demonstrate a biologically meaningful effect of the Sia residue during sperm capacitation.

## Material and Methods

### Materials

Chemicals were purchased from Sigma-Aldrich at highest research grade with the exception of the following products. Chloroform was purchased from Fronine (Riverstone, NSW, Australia). The 2-D quant kit was from G.E. Healthcare (Castle Hill, NSW, Australia). BCA assay kit was from Quantum Scientific (Pierce, Milton, QLD, Australia). HEPES was from Invitrogen Australia (Melbourne, VIC, Australia). Sequencing grade trypsin was supplied by Promega (Alexandria, NSW, Australia). Antarctic phosphatase and PNGase F were purchased from New England Biolabs (Arundel, QLD, Australia). Phospholipase A_1_ selective substrate (PED-A_1_) (N-((6-(2,4-DNP)Amino)Hexanoyl)-1-(BODIPY® FL C5)-2-Hexyl-Sn-Glycero-3-Phosphoethanolamine) was purchased from Molecular Probes (Melbourne, VIC, Australia). The TiO_2_ was collected from a disassembled column. Ionophore A23187 was purchased from Calbiochem (EMD Biosciences, La Jolla, USA).

### Sperm collection and in vitro capacitation

Animal use was approved by institutional and New South Wales State Government ethics committees. Adult Swiss mice (~8-10 weeks) were euthanized and the epididymides were removed. Sperm cells were recovered from the cauda of the epididymides using retrograde flushing [29, 30] and then incubated for 10 min at 37°C in 0.3% BSA BWW media to allow cell dispersion [31]. For the non-capacitated group, sodium bicarbonate was replaced by sodium chloride in the BWW media. Pentoxifylline and dibutyryl-cAMP (dbcAMP) were added only in the capacitated group at a final concentration of 1mM each. All samples were incubated for 60 min at 37°C and then sperm cells were washed three times (300 × g, 3 min) using BWW media without BSA.

### Protein extraction and sialoglycopeptide enrichment

Sperm pellets were resuspended in a lysis buffer consisting of 1% (w/v) C7BzO [3-(4-Heptyl) phenyl-3-hydroxypropyl) dimethylammoniopropanesulfonate], 7 M urea, 2 M thiourea, and 40 mM Tris (pH 10.4) at a final concentration of ~2.5 × 10^6^/ 100 μL and incubated for 1 h (4°C) with constant rotation. Supernatant (18,000 × g, 15 min, 4°C) was recovered and total protein was quantified using a 2-D quant kit following manufacture’s protocol. Proteins were reduced (10 mM DTT, 30 min, 30°C), alkylated (45 mM iodoacetamide, 30 min, 30°C) and 250 μg of protein was precipitated using methanol and chloroform [32]. Samples were incubated overnight (37°C) with trypsin at a 1:50 (trypsin/protein) ratio. Proteases were inactivated (bath sonication, 15 minutes) and peptides were treated with alkaline phosphatase (20 U, 2 hours, 30°C).

Enrichment of glycopeptides containing terminal Sia was performed as previously described [33]. In brief, peptide samples were diluted in loading buffer [1 M glycolic acid, 80% (v/v) ACN, 5% (v/v) TFA] and then applied to TiO_2_ beads (2 mg). After incubation for 1 h, TiO_2_ beads were washed [washing buffer 1; 80% (v/v) ACN, 1% (v/v) TFA, and washing buffer 2; 20% (v/v) ACN, 0.1% (v/v) TFA] and dried in a vacuum concentrator. Enzymatic deglycosylation of N-linked sialoglycopeptides was performed with 1 μL of PNGase F for 3h at 37°C. Released peptides were recovered, dried, resuspended in 0.1% (v/v) TFA and then loaded on a LC-MS (AmaZon ETD Ion Trap; Bruker Daltonik, Bremen, Germany) with an online-nanosprayer, and run as previously described [34].

### Bioinformatics

Acquired CID spectra were processed in DataAnalysis 4.0; deconvoluted spectra were further analyzed with BioTools 3.2 software and submitted to Mascot database search (Mascot 2.2.04, Swissprot database (546439 sequences; 194445396 residues, release date 19/20/14)). The following variable modifications have been used: phosphorylation (STY), carbamidomethylation (C), deamidation (NQ) and oxidation (M). To identify N-linked glycosylation sites, a deamidated Asn residue had to be flanked by the glycosylation consensus motif (NXS/T, where × is any amino acid besides proline) which was manually validated. Peptides that were assigned a deamidation event based solely on the MS data (i.e., no *y*- or *b*- fragment ion for a particular deamidated Asn residue could be detected) were presumed to be glycosylated only if a canonical N-glycan motif was present.

The derived mass spectrometry datasets on the 3D-trap system were combined into protein compilations using the ProteinExtractor functionality of Proteinscape 2.1.0 573 (Bruker Daltonics, Bremen, Germany). In order to exclude false positive identifications, peptides with Mascot scores below 40 were rejected. Peptides with a mascot score above 40 were manually validated in BioTools (Bruker Daltonics, Bremen, Germany) on a residue-by residue basis using the raw data to ensure accuracy as previously described [33]

### Peptide label-free quantification

MS-based label-free quantification of the N-glycopeptides identified was performed using the software Data Analysis 4.1 (Bruker Daltonik GmbH, Bremen, Germany). Peptides were matched based on charge state, m/z value and elution time. The match was confirmed by visual inspection of the peptide on the survey view and by manual comparison of the MS/MS spectra if available. Relative peptide quantification was carried out by integrating the area of the extracted ion chromatograms (XIC) of the monoisotopic peak from MS spectra (Fig. 1) [33]. For peptide quantification, four biological replicates, each consisting of pooled samples from 3-4 mice, were run.

**Figure 1.**
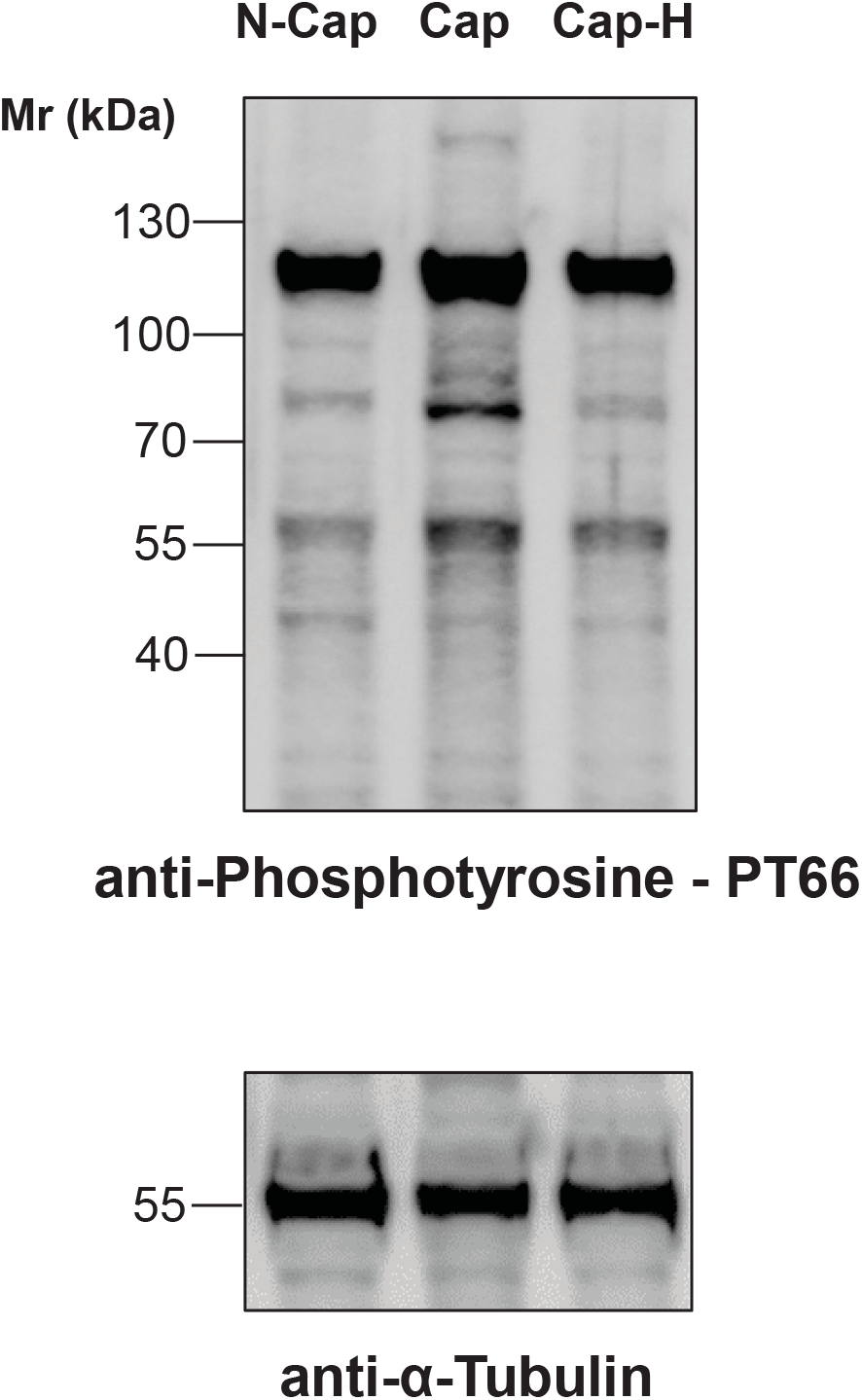
Immunoblotting of mouse cauda sperm previously incubated in non-capacitating (N-Cap) or capacitating media supplemented (Cap-H) or not (Cap) with the PKA inhibitor H89. Protein extraction was performed in SDS-PAGE buffer and 10 μg was loaded per lane. Membranes were probed with anti-Phosphotyrosine PT66 antibody and then re-probed with anti-a-tubulin. The data show one replicate that has been repeated in 6 biological replicates.

### PLA_1_ activity assay

The phospholipase A_1_ (PLA_1_) activity of EL in intact mouse spermatozoa was analyzed using a dye labeled-PLA_1_ specific substrate [35]. Sperm samples were divided into six groups; non-capacitated with or without H89 or A23187 and capacitated with or without H89 or A23187. The compound H89 was supplemented 10 min prior to the addition of pentoxifylline and dbcAMP. After capacitation for 1h, ionophore A23187 was added to cells (20 μM final concentration) after 30 min of incubation and then samples were allowed to incubate for another 30 min. Spermatozoa were washed (300 × g, 3 min) twice using BWW media with BSA and then once with BWW media without BSA. Sperm pellets were then resuspended in BWW media without BSA to a final concentration of 20 × 10^6^sperm/ mL.

A stock solution of PED-A_1_ substrate (5 mM in DMSO) was diluted (1:625) with BWW medium without BSA yielding a final concentration of 8 μM. Aliquots of 25 μL of the PED-A_1_ working solution were placed in a v-bottom 96-well plate and then 25 μL of the sperm samples was added. Controls included wells containing only PED-A_1_ solution or sperm cells. The plate was incubated at 37°C and fluorescence measurements were taken every 30 sec during a 1h-period using the microplate reader FLUOstar OPTIMA (BMG Labtech, Mornington, VIC, Australia) with excitation and emission wavelengths of 485 and 510 nm, respectively. Linear regression was calculated using the average results for the first 10 min of reading and reaction rates (slope) were calculated for each sample. PLA_1_ activity was assessed in five independent experiments.

#### Aconitase Vectors

ACO2 cDNA was ligated into the pcDNA3-EGFP plasmid. Primers were designed to have HindIII (forward) and NotI (reverse) restriction sites as well as 8 extra base pairs on the ends to allow enzyme cleavage. Two different reverse primers were used to create two different plasmids, the first containing a linker sequence to create an ACO2-EGFP fusion, and the second containing a linker sequence followed by a (HIS)_6_-tag and stop codon to create his-tagged ACO2. This was achieved using mouse cDNA and the following overhanging primers:

Forward: ACGAATTC–AAGCTT–ATGGCGCCTTACAGCCTCCTGGT

Reverse 1: GGTGCTTA–GCGGCCGC–GAGCTTCCACCACCTCC– CTGCTGCAGCTCCTTCATCCTGTTG

Reverse 2: GGTGCTTA–GCGGCCGC–TCA– ATGGTGGTGGTGATGATGGCTTCCACCACCTCC– CTGCTGCAGCTCCTTCATCCTGTTG

For PCR, Thermo Scientific Phusion High-Fidelity DNA Polymerase was used and their instructions followed. Each 20 μl PCR was made up with the following concentrations: 1 × HF buffer, 200 μm dNTPs, 0.5 μM each of forward and reverse primer, ~100 ng DNA, 3% DMSO and 1 unit Phusion polymerase. PCR conditions were: 98°C for 30 sec, [98°C for 10 sec, 65°C for 30 sec, 72°C for 2 min] × 35 cycles, 72°C for 10 min. Annealing temperature was optimized at 65°C for both sets of primers.

PCR inserts and pcDNA3-EGFP vector were digested with Promega enzymes; NotI and HindIII after Wizard mini-prep kit clean-up. Insert and vector were ligated at a 3:1 ratio using Promega T4 DNA ligase and then transformed into E. coli cells on ampicillin agar plates. Singles colonies were cultured the following day and then plasmids extracted. Plasmids were again digested with HindIII and NotI to check for insertions and the ones containing inserts were sent away for Sanger sequencing using four sequencing primers to cover entire insert and check for mutations: GGACTTTCCAAAATGTCG, AGGCCGAACAGACATTGC, AGATGCAGACGAGCTTCC, TTCATCCAGTGGACAAGC.

### Site directed mutagenesis

To mimic the negative charge of the Sia modification on ACO2, we changed the amino acid 612 (Asn or N) to aspartic acid (D) using site directed mutagenesis. A single base pair change on chr15:81913178 to change the codon from AAC to GAC was achieved using the following primers:

Forward: TGCTCATCGGTGCCATCAACATC

Reverse: GGTTGTTAGAGATGTCATCCAGATGCCCAC

Phusion DNA Polymerase PCR was done using the same reaction as written in the above methods and the PCR conditions as follows: 98°C for 3 min, [98°C for 10 sec, 64 °C for 30 sec, 72°C for 6 min] × 35 cycles, 72°C for 10 min. Following PCR, the plasmid bands were cut out of an agarose gel, purified and digested with DpnI from NEB. Following another clean-up step, the plasmids were transformed into E. coli and grown on agar plates overnight. Single colonies were selected the following day and cultures grown overnight. Plasmids were extracted from these colonies and sent away for sequencing to check for mutation using sequencing primer AGATGCAGACGAGCTTCC.

### Transfection

HEK293T cells were used for transfection with ACO2-EGFP plasmid and ACO2-(HIS)6 plasmid including the WT and N612D versions of these plasmids. We confirmed that human cells would be suitable for transfection with the mouse ACO2 gene due to almost identical sequences. Cells were split into 6-well plates to attain ~50% confluence the following day. Transfection was done with 5 μg plasmid DNA and 10 μg PEI in 3 mL media. Firstly, 5 μg of plasmid and 10 μg PEI were put into separate tubes and 150 μL DMEM (no FBS) added to each one. These were then mixed together and incubated for 30 min. Next, media in the 6-well plates was replaced with 2.7 mL of fresh DMEM and the 300 μL DNA:PEI mix added. Cells were left to transfect for the times indicated and then harvested. ACO2-GFP cells were fixed using 4% paraformaldehyde for microscopy and FACS analysis or frozen at −80°C for immunoblotting. ACO2-(HIS)_6_ cells were frozen at −80°C for immunoblotting or Aconitase assay.

### Aconitase Assay

#### Aconitase assay from BioVision was used with adjusted methods

Day 1: Frozen transfected HEK293T cells consisting of (HIS)_6_-wild-type ACO2 transfected (WT) and N612D mutant and control non-transfected cells were thawed on ice. Cells were resuspended in 400 μL lysis buffer (PBS pH 8, 10 mM imidazole, 0.5% tween) and then sonicated and centrifuged (16,000 g, 4°C) for 15 min. A 200 μL aliquot of N612D and control samples were placed in fresh tubes and the rest discarded. The WT samples were split into 3 tubes: 200 μL, 100 μL and 50 μL. A 200 μL of lysis buffer and then 50 μL Ni-NTA agarose beads were added and samples rolled for 1 h at 4°C. Beads were washed thrice in wash buffer (PBS pH 8, 20 mM imidazole, 0.5% tween) and once in kit assay buffer. Afterwards, beads were resuspended in 100 μL kit assay buffer, 10 μL activation solution was added, and samples were rolled for 1 h at 4°C. One hundred μL of reaction mix was added to each tube and samples rolled at room temperature overnight.

Day 2: Frozen non- and capacitated sperm cells were thawed on ice. Cells were resuspended in 100 μL kit assay buffer, sonicated and centrifuged (16,000 g; 4°C) for 15 min. Ten μL of activation solution was added and samples incubated for 1 h on ice. Afterwards, 100 μL reaction mix was added to each tube and samples incubated at room temperature for 2.5 h. Standards were made according to instructions and allowed to incubate for 30 min. The beads from day-1 preparation were centrifuged and, together with the non- and capacitated sperm prepared on day 2, were loaded in a 96-well plate in 100 μL duplicates. Ten μL of developer was added, samples incubated at room temperature 25°C and read at 450 nm.

Following kit, nickel bead enrichment, (HIS)_6_-recombinant protein were eluted with 250 mM imidazole and then frozen along with non- and capacitated samples. For immunoblotting, samples were methanol/chloroform precipitated and lysed in SDS-PAGE buffer.

### SDS-PAGE and immunoblotting

Non-capacitated and capacitated mouse spermatozoa (with and without H89) were prepared as described above and then diluted in SDS-PAGE buffer. Protein (10 μg) was separated by SDS-PAGE using 4-20% precast polyacrylamide gels (NuSep Ltd, Lane Cove, NSW, Australia) and then transferred onto nitrocellulose membrane Watman® Optitran® BA-S 85 (GE Healthcare, Castle Hill, NSW, Australia). The membrane was blocked (1 h at room temperature) and incubated overnight at 4°C with rabbit polyclonal antibody raised against EL (orb100394, LIPG; Biorbyt) at a dilution of 1:500 in 5% (w/v) skim milk TBS-T (0.02 M Tris, 0.15 M NaCl, 0.1% (v/v) Tween-20; pH 7.6).

After three washes, membrane was incubated for 1 h at room temperature with anti-rabbit IgG horseradish peroxidase (HRP) conjugate (Sigma-Aldrich) at a concentration of 1:1000 in 5% (w/v) skim milk TBS-T. The membrane was washed thrice, and immuno-reacted proteins were detected using an enhanced chemiluminescence (ECL) kit (Amersham International) according to the manufacturer’s instructions. Equal loading was confirmed by stripping the membrane and then re-probing it with a mouse monoclonal anti-a-tubulin antibody.

To confirm the capacitation of sperm cells and the efficiency of H89 at inhibiting tyrosine phosphorylation via PKA, samples were probed with the mouse monoclonal anti-phosphotyrosine–peroxidase antibody PT66 (A5964; Sigma-Aldrich). Loading control was performed using mouse monoclonal anti-a-tubulin antibody.

Immunoblots of WT and mutant ACO2 were essentially performed as previously described [34]. The following antibody dilutions were used: rabbit polyclonal anti-Aconitase 2 antibody at 1:1000 (ab83528; Abcam), rabbit polyclonal anti-6X His tag antibody at 1:1000 (ab1187; Abcam). All secondary antibodies were used at a 1:1000 dilution. Anti-GFP was a kind gift from Peter Lewis and used at 1:1000.

### Immunocytochemistry

Capacitated and non-capacitated intact mouse spermatozoa were fixed in 4% (w/v) formaldehyde for 10 min at room temperature. Cells were washed three times in PBS containing 0.5 M glycine, immobilized on poly-L-lysine coverslips and permeabilized with ice cold methanol for 10 min. Coverslips were washed, blocked for 1 h with 3% (w/v) BSA in PBS and incubated overnight with anti-EL antibody in a 1:50 dilution with PBS containing 1% (w/v) BSA. Following three washes with PBS, cells were incubated with Alexa Fluor® 488 goat anti-rabbit IgG (Life technologies) in a 1:100 dilution with PBS containing 1% (w/v) BSA. Coverslips were washed as described above and then mounted with Mowiol antifade medium. Cells were evaluated using phase contrast and epifluorescence microscopy.

### Acrosomal status

The acrosomal status of mouse spermatozoa was assessed either at time 0 (non-capacitated) or following 30 min of capacitation (37°C; 5% CO2). After 30 min of incubation, the ionophore A23187 (10 μM final concentration in DMSO) or DMSO vehicle only were added and then cells were incubated for 30 min. Samples were washed twice using PBS and 10 μl of sperm suspension was spotted onto superfrost slides, spread with a glass pipette and air-dried. Slides were immersed in absolute methanol for 15 min, rinsed with PBS and then incubated for 30 min with fluorescein isothiocyanate-conjugated *Arachis hypogaea* peanut agglutinin (FITC-PNA) (15 μg/μL final concentration). The slides were rinsed in PBS and mounted with antifade media. For each slide, images of adjacent fields were recorded under a x40 magnification to achieve a sperm count of at least 100 spermatozoa. Spermatozoa were classified into one of the three categories of FITC-PNA labelling: I = intact acrosome; II = partial acrosome reaction; and III = complete acrosome reaction. For each experimental condition, a minimum of 3 slides were examined and quantified.

### Statistical analysis

The data obtained by MS-based label-free quantification were normalized among runs using the average area of five different glycopeptides visually selected based on their quality and constant intensity. The normalized area of each glycopeptide was then compared between non-capacitated and capacitated sperm samples using student’s *t*-test. Relative immunoreactivity for EL and a-tubulin and sperm PLA_1_ activity were compared among groups using paired *t*-test. P-values < 0.01 were considered as significant. Standard errors are shown in the graphs. The data for acrosome reaction from 5 biological replicates were subjected to analysis of variance (two-way ANOVA) followed by paired *t*-test. Data are shown as mean ± S.D. when not specified otherwise.

### Molecular modelling; Covalent docking model of Sialic acid binding of Asn612 to aconitate hydratase

Molecular docking experiments were carried out using the program GOLD (Genetic Optimization for Ligand Docking) version 5.2 and favouring the CHEMPLP Scoring system (Verdonk 2003). The three-dimensional iteration of Sialic acid (MolPort-008-267-866) was used for covalent docking onto the identified target residue of Asn612 on a model of Aconitate hydratase, mitochondrial precursor from *Mus musculus* (EC: 4.2.1.3) generated by the PHYRE2 protein recognition server Kelley (2015). The proposed reaction mechanism and subsequent product is depicted in Scheme 1. The size of the search domain was set to 10 Å and the covalent docking function within GOLD was employed. Docking was executed using a 100% search efficiency, generating ten Genetic Algorithm (GA) runs, while the rotameric states of several sidechains including Gln563, Lys605 and Arg648 were set to library rotamer orientations and the remainder of the site remained rigid. The generated binding poses were then inspected, and conformations were chosen for further analysis taking into account their ranking and interactions with the probed residues. Molecular visualizations were performed using the software package PYMOL (Schrödinger, NY, USA).

## Results

In order to determine the efficiency of the protocol used here to induce sperm capacitation, tyrosine phosphorylation was measured (Fig. 1). As shown, tyrosine phosphorylation increased during capacitation, which could be abrogate with the PKA inhibitor H89, suggesting that the sperm cells used in this study were capacitating. The experimental paradigm used in this study was to compare the changes associated with the capacitation process itself (Cap *vs* Cap-H, Fig. 2) as opposed to changes over time (N-Cap *vs* Cap, Fig. 2).

**Figure 2.**
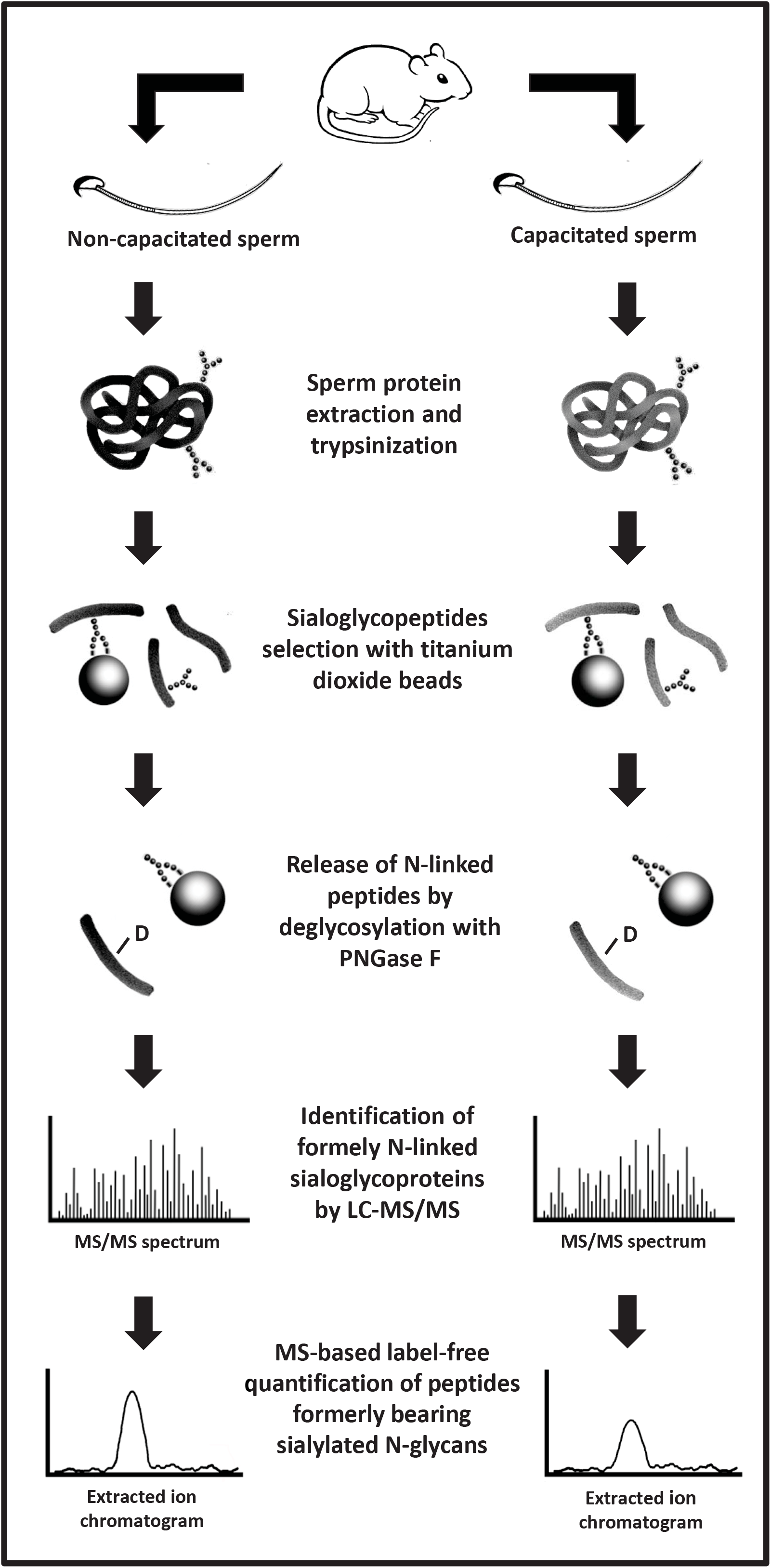
Schematic representation of the strategy used to identify and quantify sialylated N-linked glycopeptides extracted from mouse sperm. Deamidated asparagine residue (-D).

### Sialylated glycoproteins identified in mouse sperm

The protocol to enrich for sialylated N-linked glycopeptides is shown in Figure 2. Herein, spermatozoa were obtained from the mouse epididymides, combined, and then separated into two separate tubes. Whilst one tube had capacitating media, the other lacked sodium bicarbonate which is essential for this process. The samples were washed, lysed and digested. Sialylated-glycoprotein enrichment was performed using TiO_2_ beads. Although this is not specific for glycopeptides, but rather for negatively charged molecules, the elution through PNGase F treatment of the beads allows the selection of N-linked glycopeptides generally or sialylated glycopeptides specifically. Enrichment of sialylated N-linked glycopeptides using TiO_2_ allowed the identification of 142 unique peptides, which were from 90 different glycoproteins (Table 1), demonstrating that some proteins possessed N-linked glycopeptides with terminal Sia in more than one position. Of particular interest, the characterization of the glycopeptides containing Sia residues represented 36.7% of the total non-reductant peptides identified by MS/MS. This is in perfect agreement with our previous paper with rat sperm, in which around 34% of the peptides identified were classified as glycopeptides after enrichment using TiO_2_ [33].

**Table 1.**
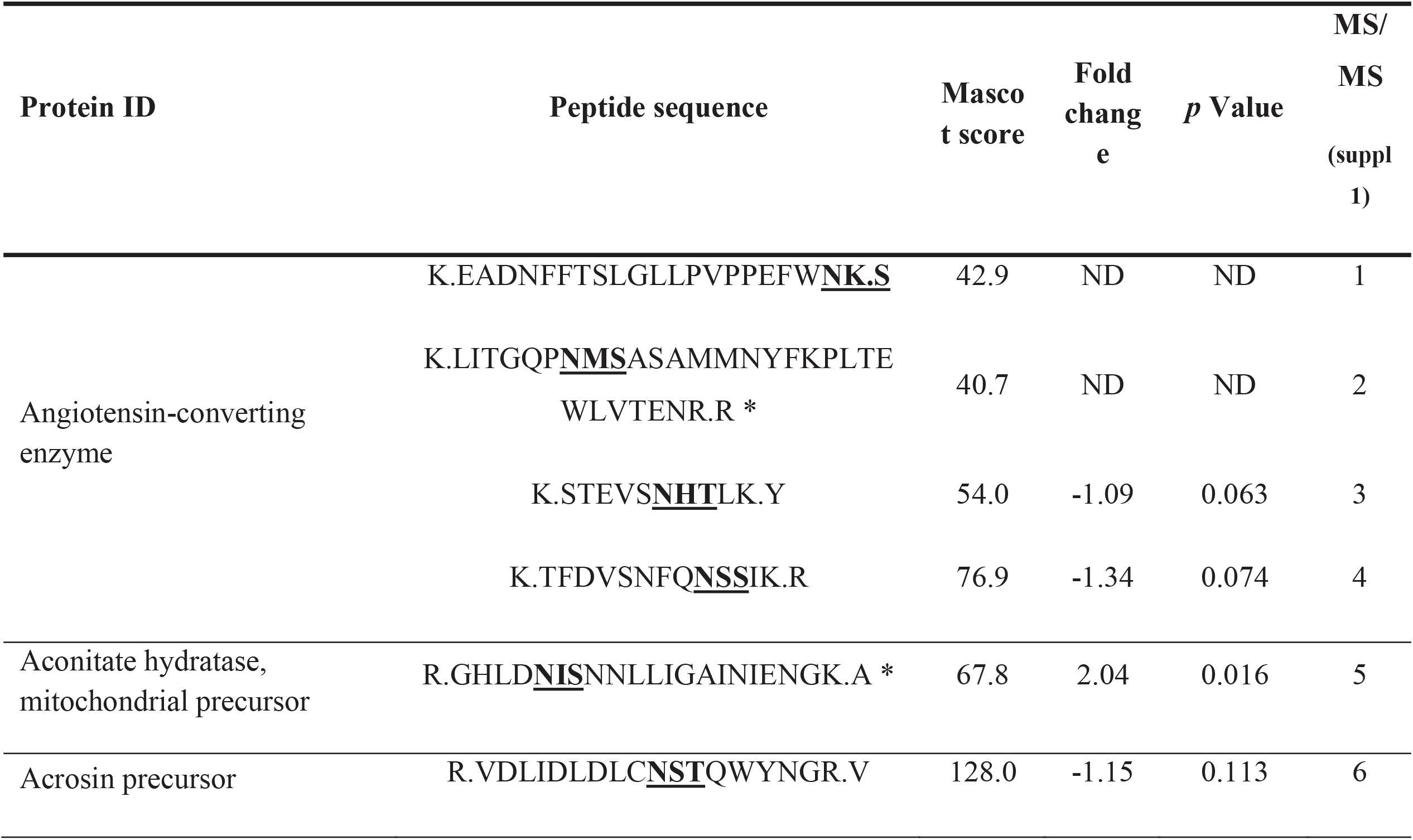
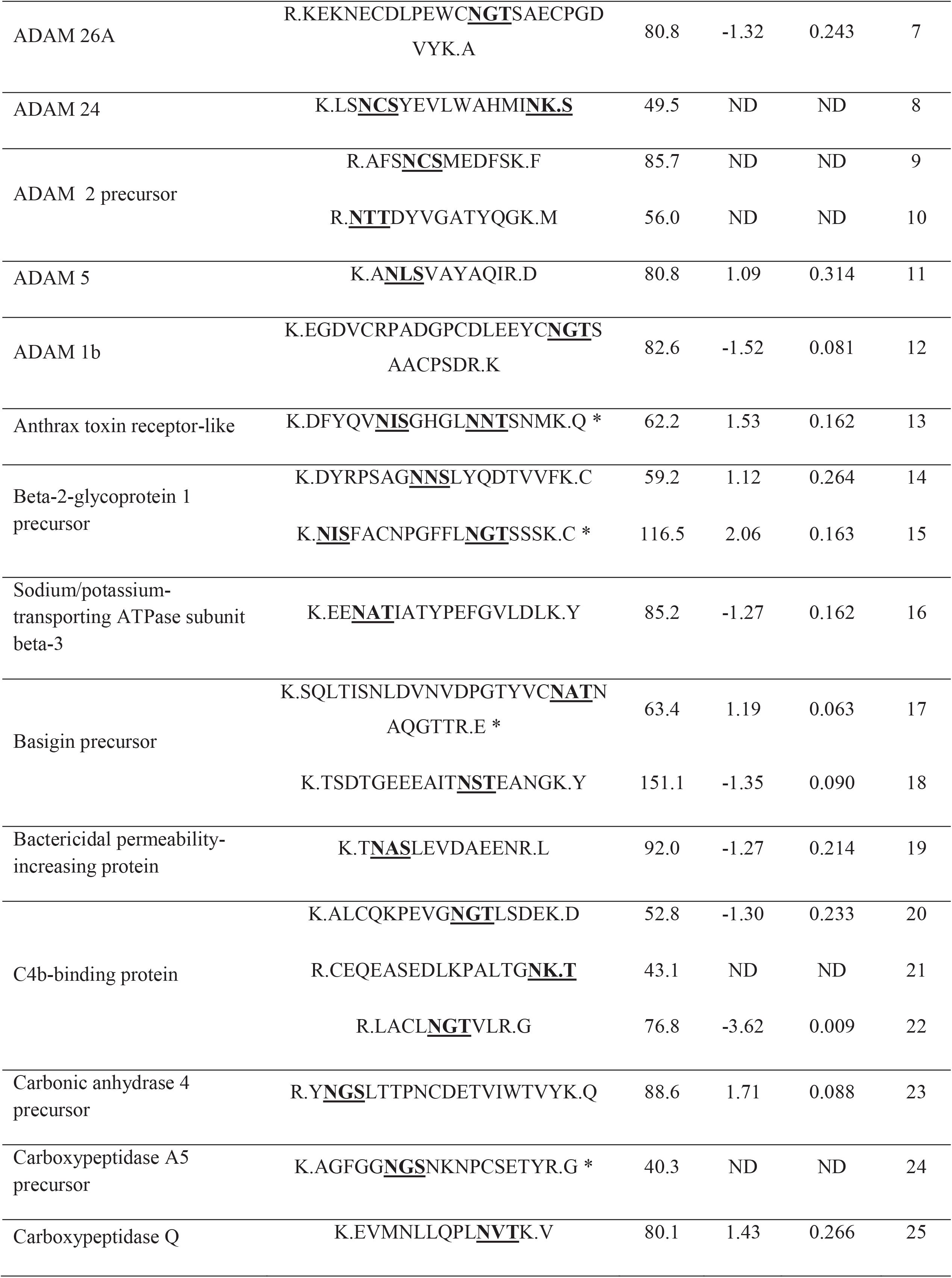
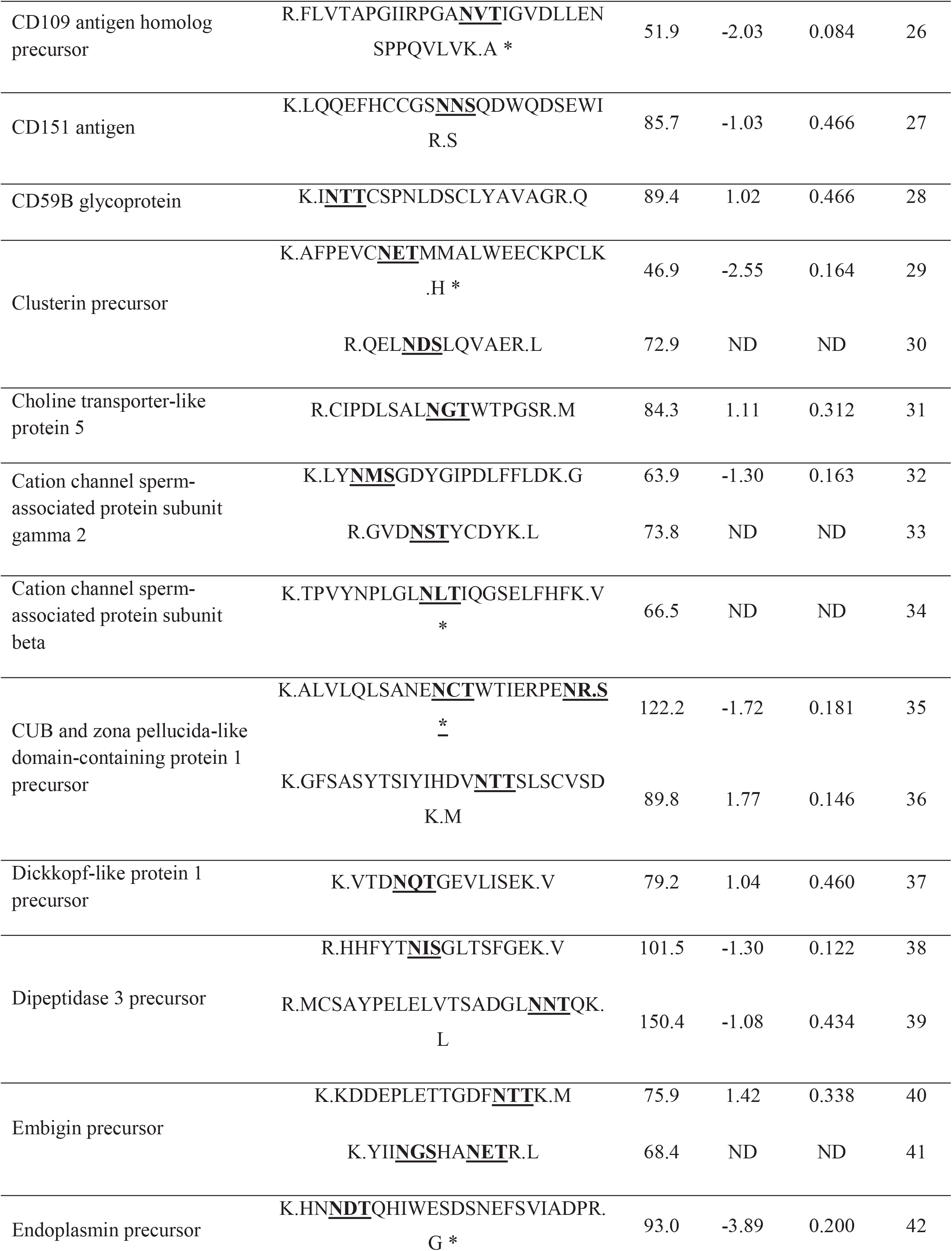
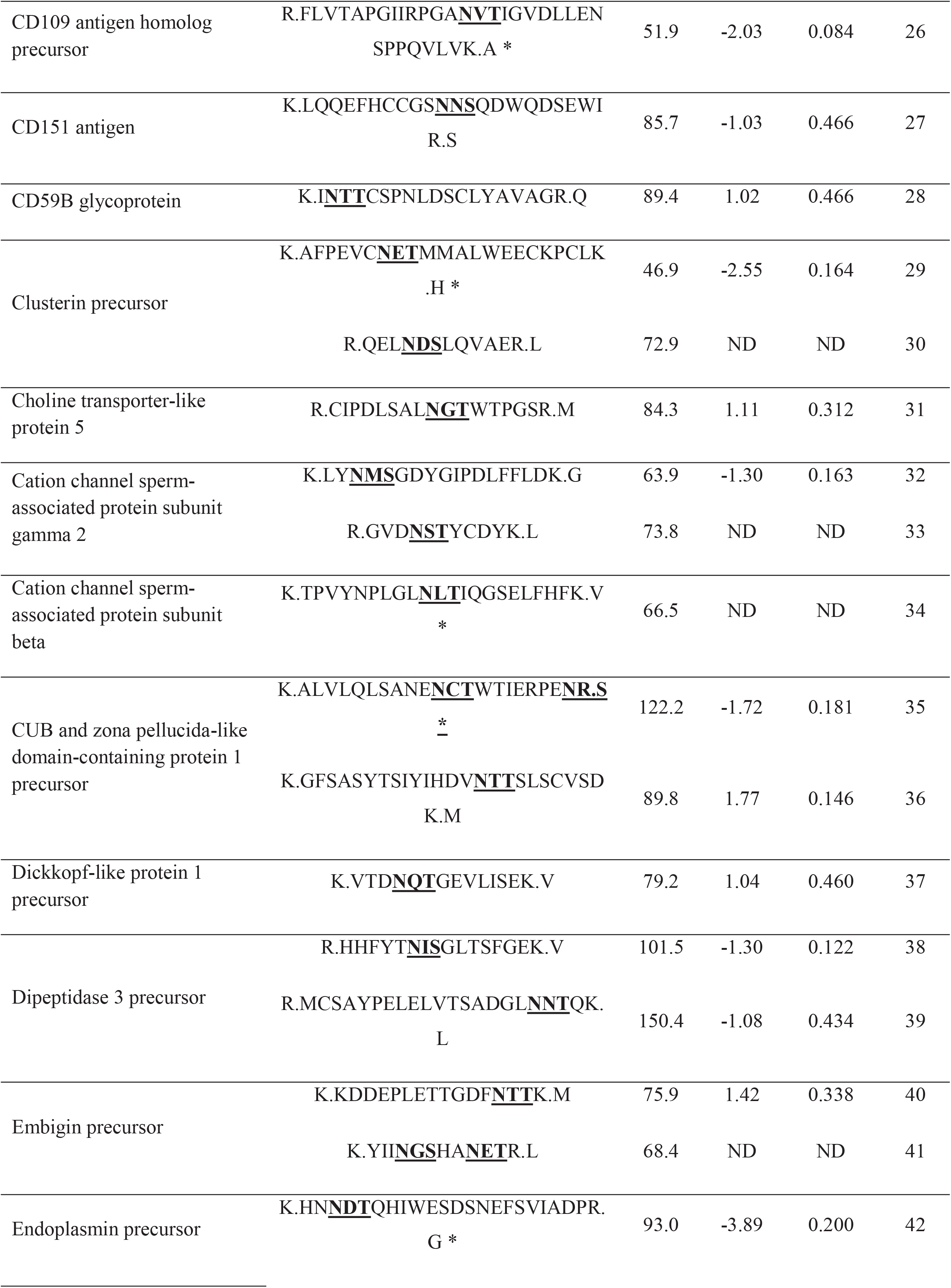
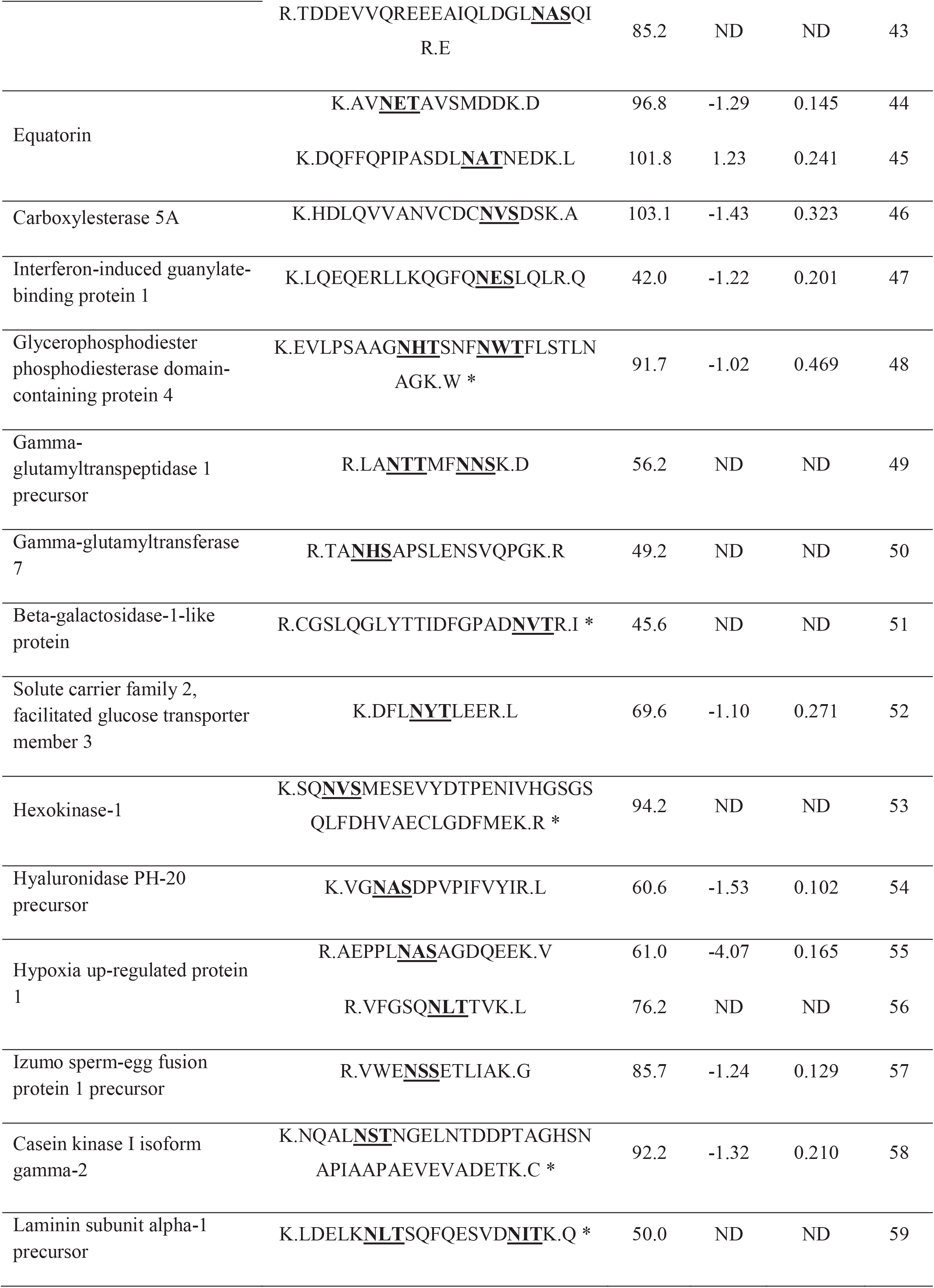
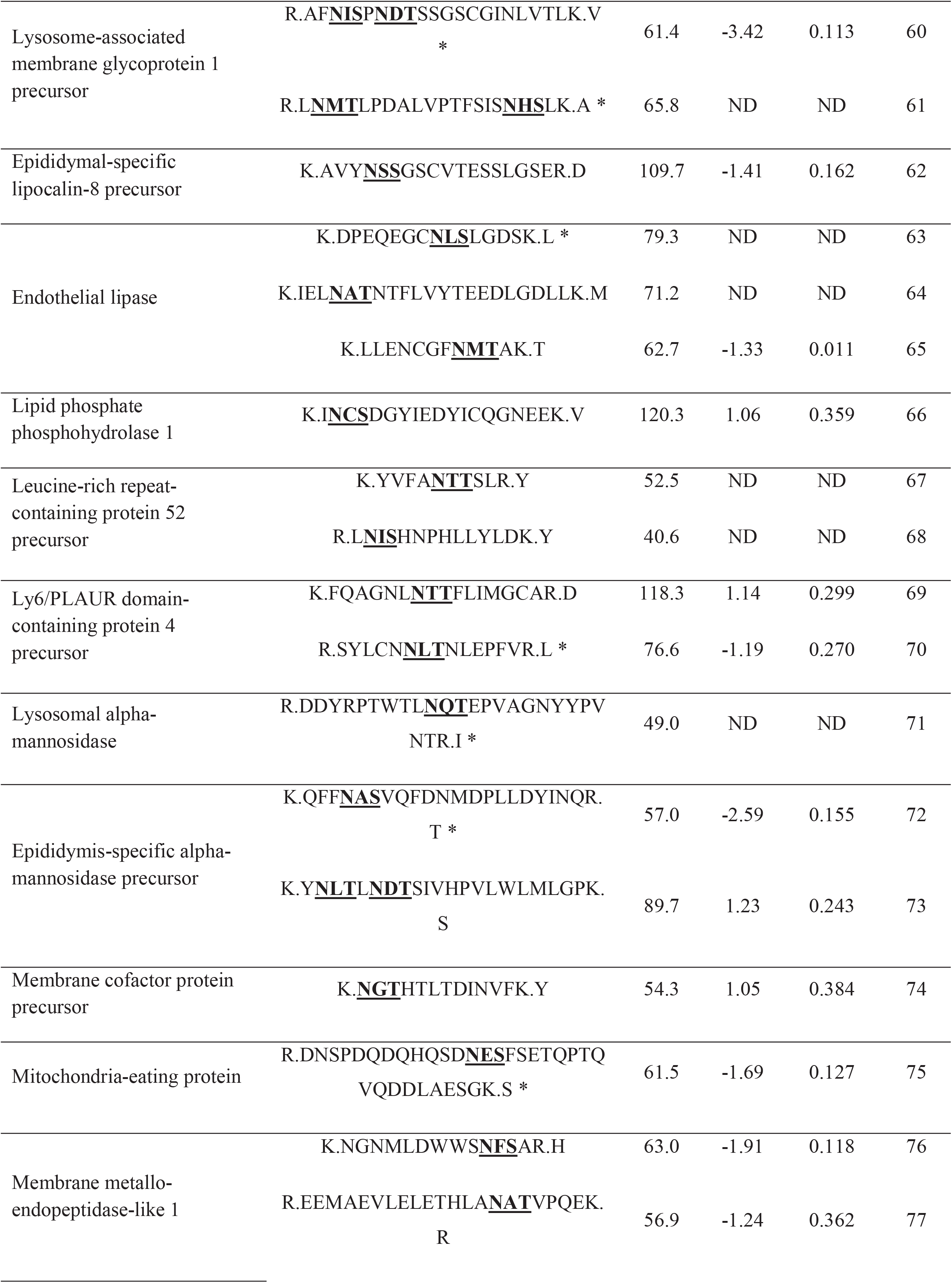
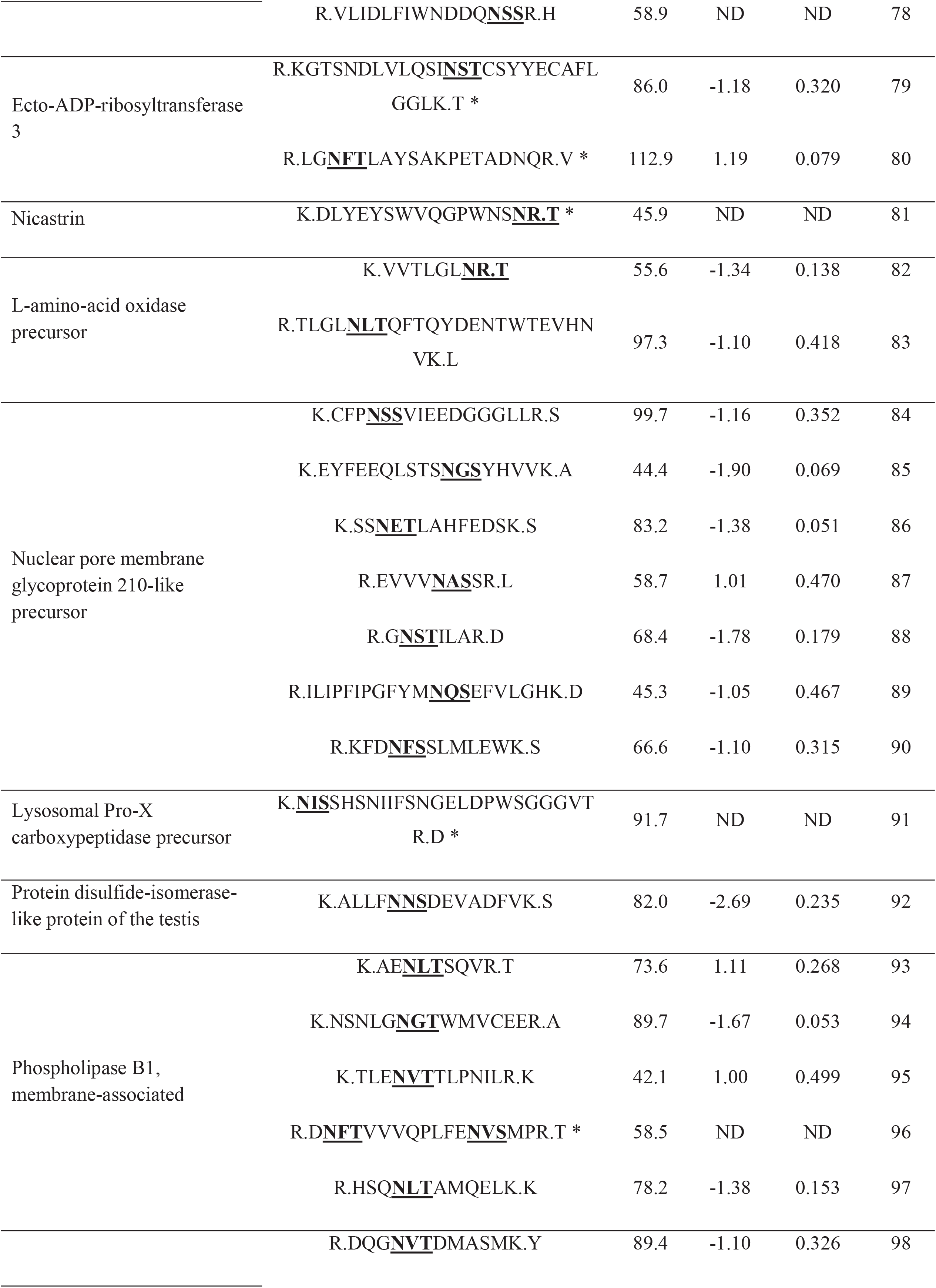
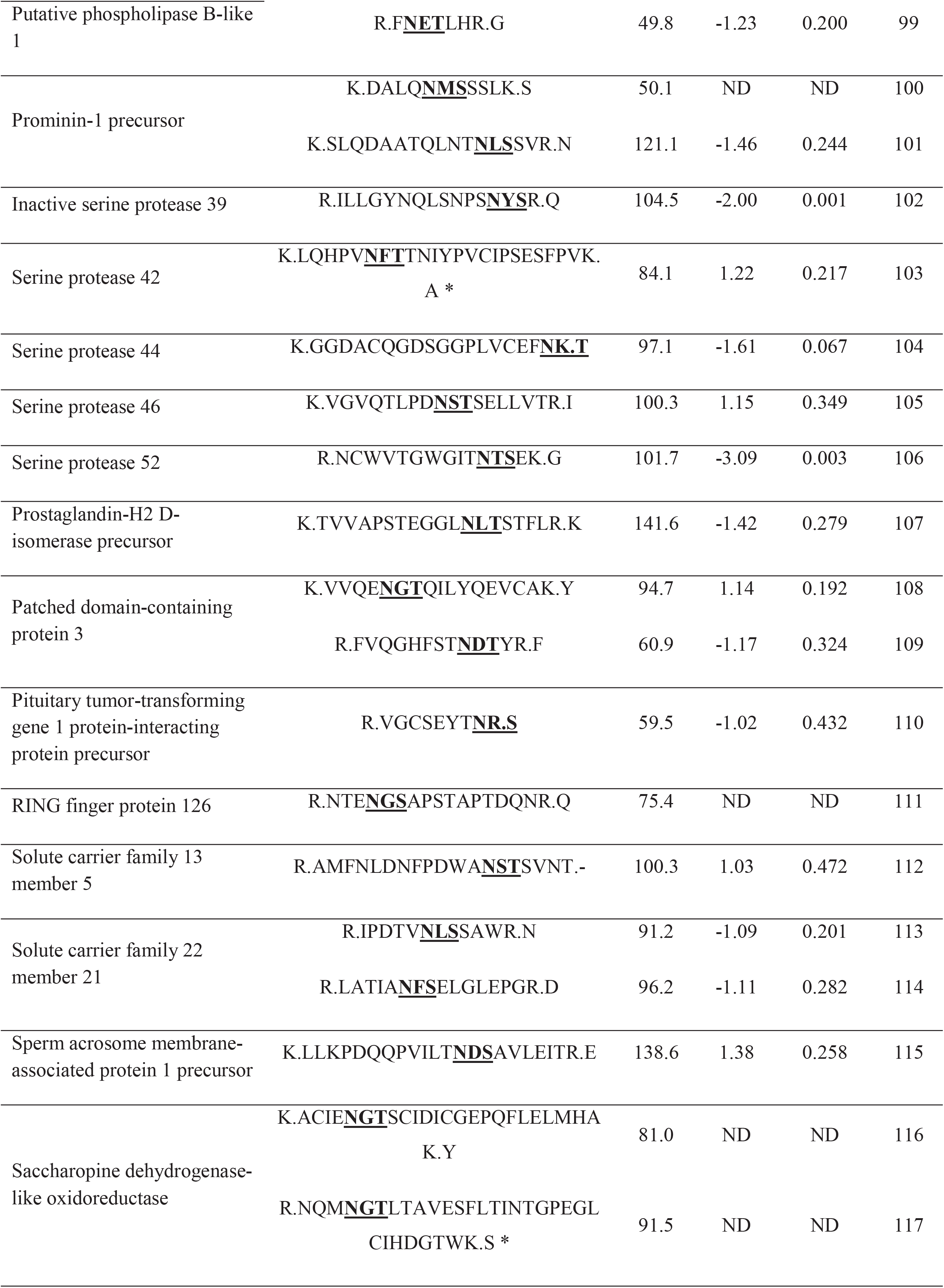
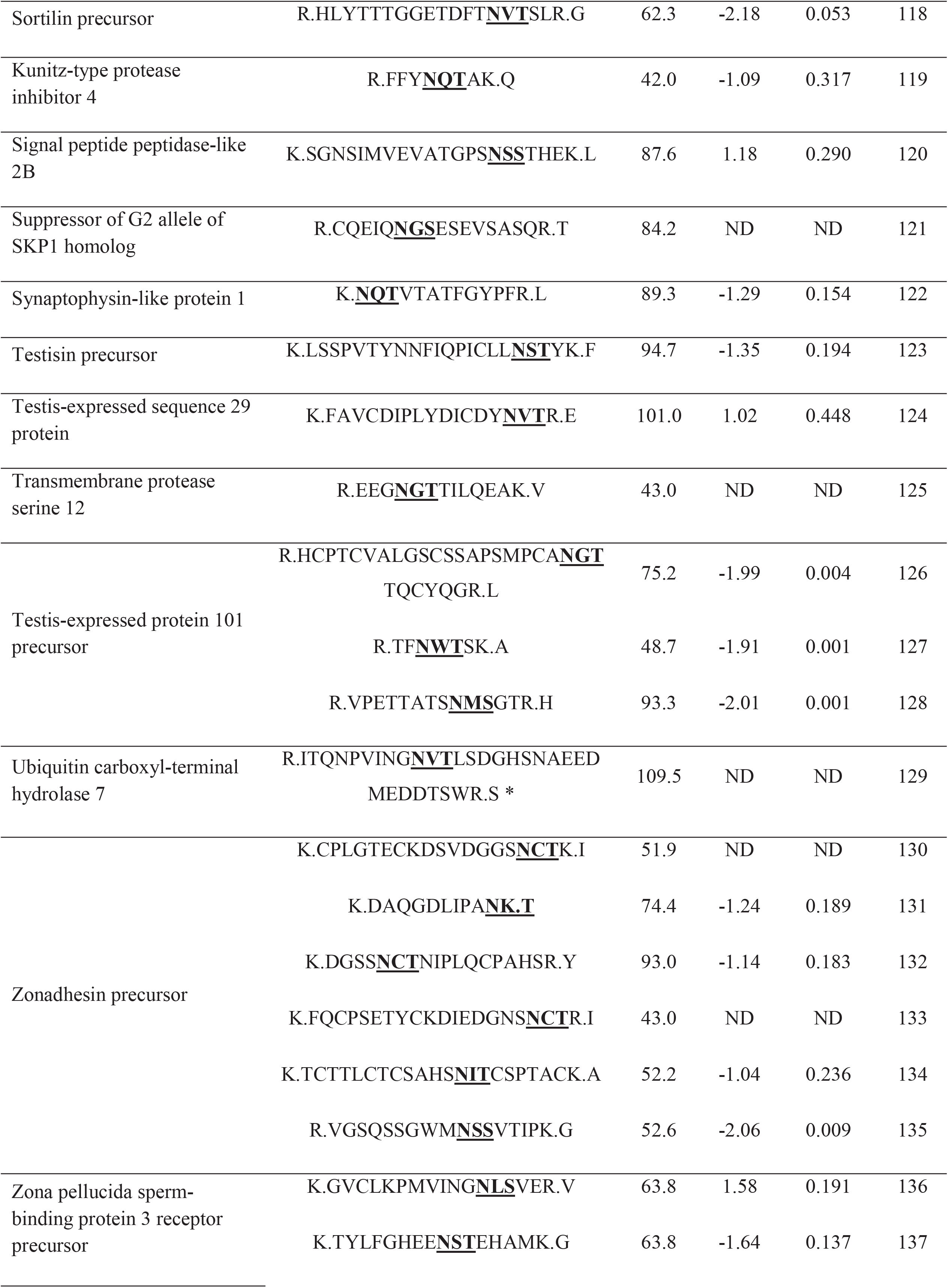
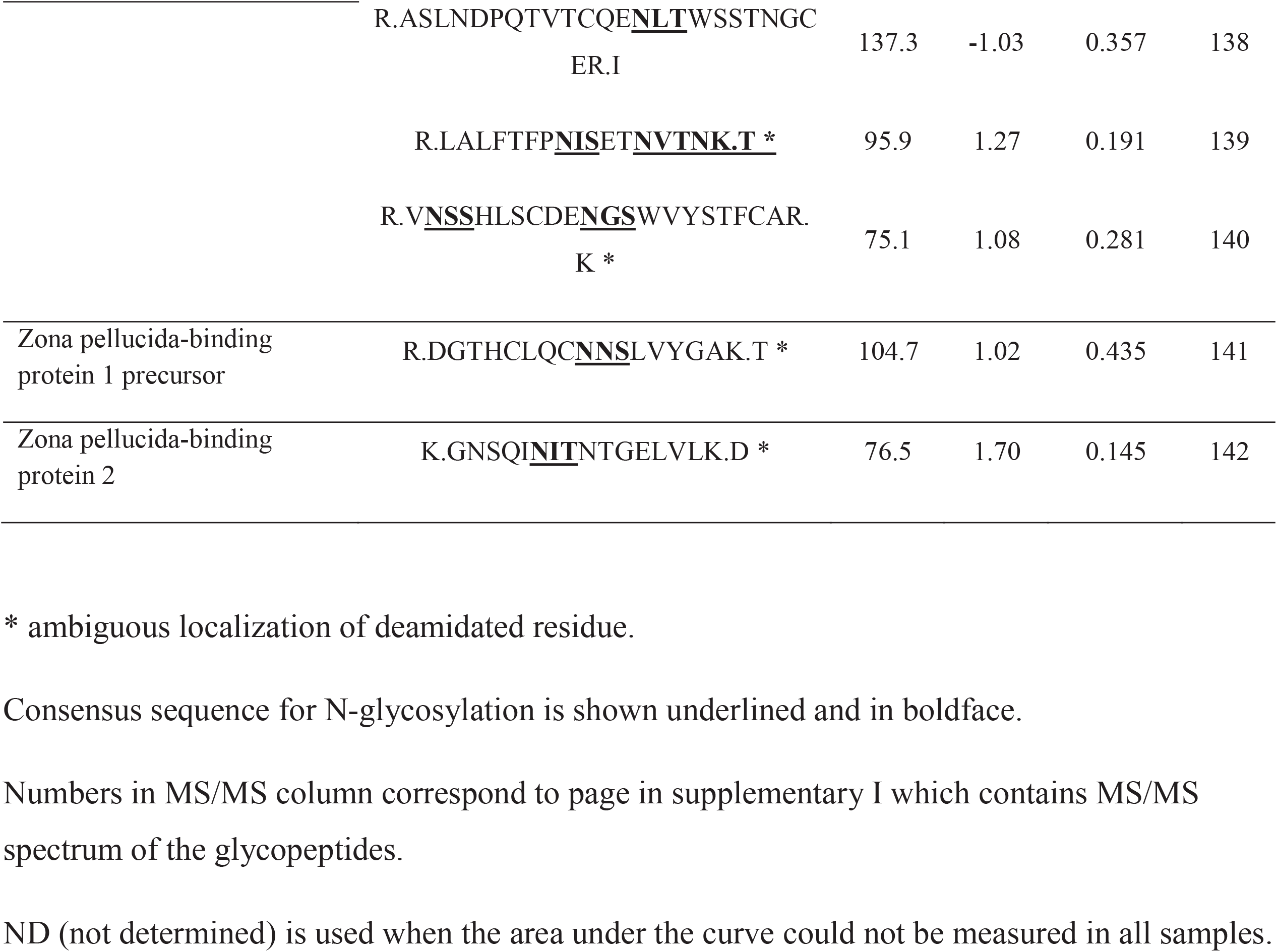
List of sialylated N-glycopeptides identified in non-capacitated and capacitated mouse sperm and their regulation during the process of capacitation.

Due to ion fragmentation patterns (i.e., no visualization of the *y*- or *b*- fragment ion for the deamidated Asn), a precise identification of the residue containing the deamidation was not possible in all cases. Hence, the peptides exhibiting ambiguous annotation of the deamidated residue are indicated in Table 1 with an asterisk. Supplementary I contains MS/MS spectrum from all peptides present in Table 1 whereas Supplementary II presents data such as measured m/z values, peptide charge and retention times.

### Changes in sialylated N-linked glycopeptides during capacitation

The data for label-free quantification of the N-linked glycopeptides in non-capacitated versus capacitated sperm are shown in Table 1. Some glycopeptides were presented at very low amount, impairing their quantification in some biological replicates. These cases are indicated in Table 1 as not determined (ND) since we were not confident to report these further. Of interest, nine (6.3%) of the glycopeptides identified here underwent significant changes during sperm capacitation. These glycopeptides belong to the proteins: ACO2, C4b-binding protein (C4BP), EL, inactive serine protease 39 (PRSS39, also known as testicular-specific serine protease 1; TESP1), serine protease 52 (PRSS52, also known as testicular-specific serine protease 3; TESP3), testis-expressed protein 101 (TEX101) and zonadhesin (Table 1). Interesting, these glycopeptides were shown to be reduced during capacitation except for the peptide from ACO2, a protein involved in the tricarboxylic acid (TCA) cycle (Table 1).

### Determination of EL amount and sub-cellular localization

In order to determine the consequences of sialylation changes during capacitation, we choose two proteins for further study: the EL and ACO2. The EL has been reported as primarily having PLA_1_ activity [36]. This was of particular interest, since changes in the composition of lipids from cell membranes, such as the formation of lysophospholipids by phospholipids hydrolysis, are known to modify membrane fluidity [37, 38] and function through the modulation of receptors [39], channels [40, 41] and enzyme activity [42] within its structure. Considering that morphological and functional changes in sperm membranes, including alteration in membrane fluidity, are essential for capacitation [43] and that lysophospholipids may be involved in the modulation of the acrosome reaction [44], we further investigated the behavior of EL during this process.

Immunoblotting for EL was performed, aiming to determine what happens to the entire protein during capacitation. As shown in Figure 3, the total amount and the molecular weight of EL did not change after capacitation (Fig. 3A, lanes 1 and 2). In addition, the use of the PKA inhibitor H89 during capacitation also did not affect the amount of EL (Fig. 3A, lanes 2 and 3). To demonstrate equal loading, we re-probed the sample with anti-α tubulin antibody (Fig. 3B). Using the software Image J, the quantitative values of each band were plotted to confirm that no significant change occur in EL expression after capacitation (Fig. 3C).

**Figure 3.**
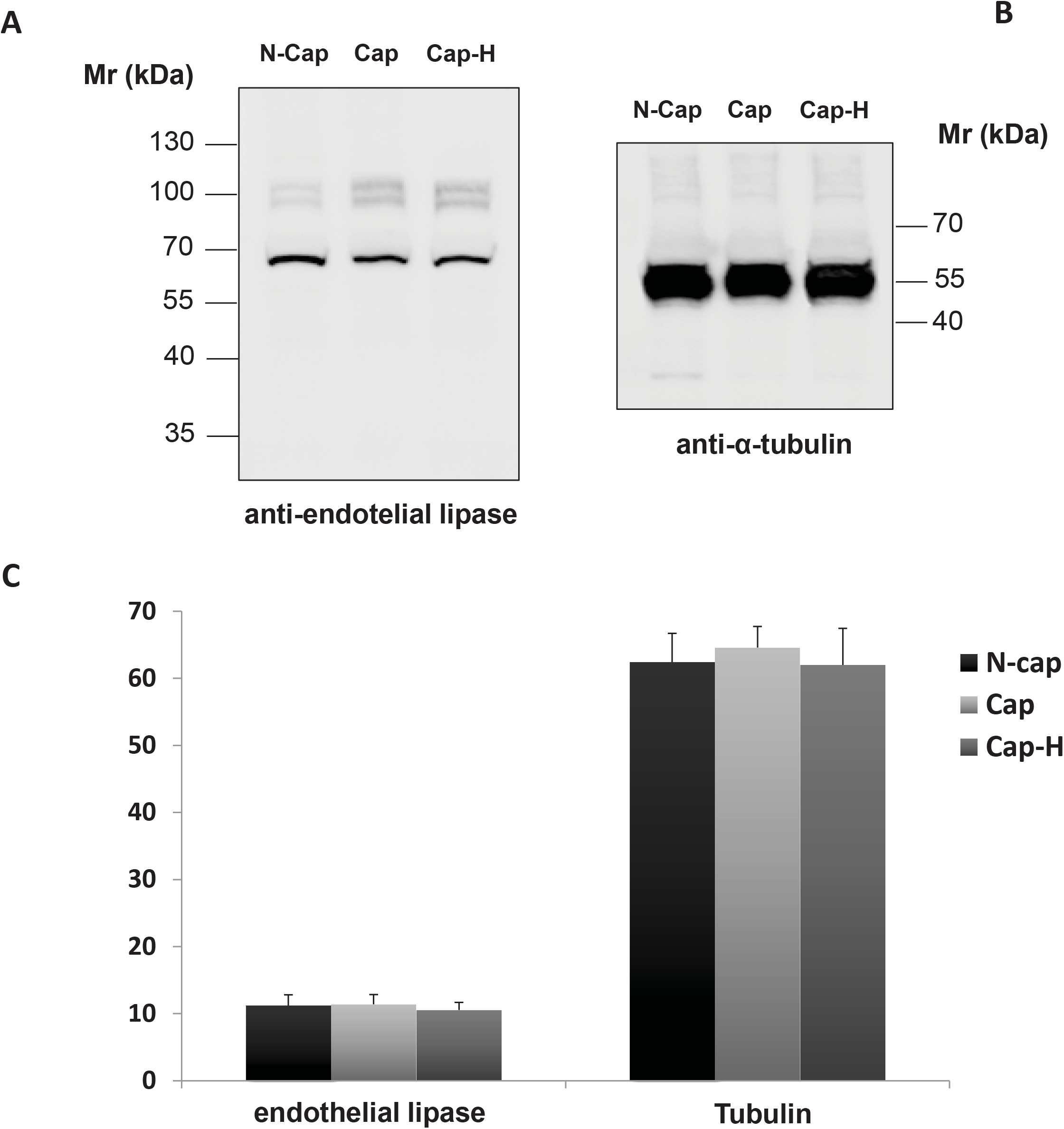
Immunoblot analysis of cauda spermatozoa from mouse. Sperm samples were incubated in non-capacitating (N-Cap) or capacitating media supplemented (Cap-H) or not (Cap) with the PKA inhibitor H89. Extraction was performed using SDS-PAGE buffer and 15 μg of protein was loaded per lane. **(A)** Membranes were probed with anti-EL and then **(B)** re-probed with anti-a-tubulin. **(C)** Graphic below shows relative immunoreactivity levels of both EL (left) and a-tubulin (right). No statistical difference was observed among groups, which consist of five biological replicates each.

To verify whether the protein EL is redistributed in spermatozoa during capacitation, we performed immunostaining using anti-EL antibody. Immunostaining for EL was observed in both head and tail regions of non-capacitated (Fig. 4a,c) and capacitated (Fig. 4b,d) mouse sperm. For both groups, the anterior acrosome region stained for EL whereas the equatorial region showed no staining. In addition, the staining intensity of the postacrosomal sheath showed high variation among cells; being absent in some cases (Fig. 4b, white arrow exemplifies this variation). In the tail region, although the midpiece and the cytoplasmic droplet showed high staining for EL, we noted that some cells exhibited weaker labeling of the midpiece (Fig. 4c,d). Of interest, the percentage of these cells, with reduced immunoreactivity for EL at the midpiece, increased when sperm was incubated in capacitating conditions (from around 7% to 68%). The same was observed in samples capacitated with the PKA inhibitor H89, suggesting that the diminish in EL immunofluorescence within the midpiece is independent of PKA (data not shown). Secondary only controls showed no fluorescence (Fig. 4e,f).

**Figure 4.**
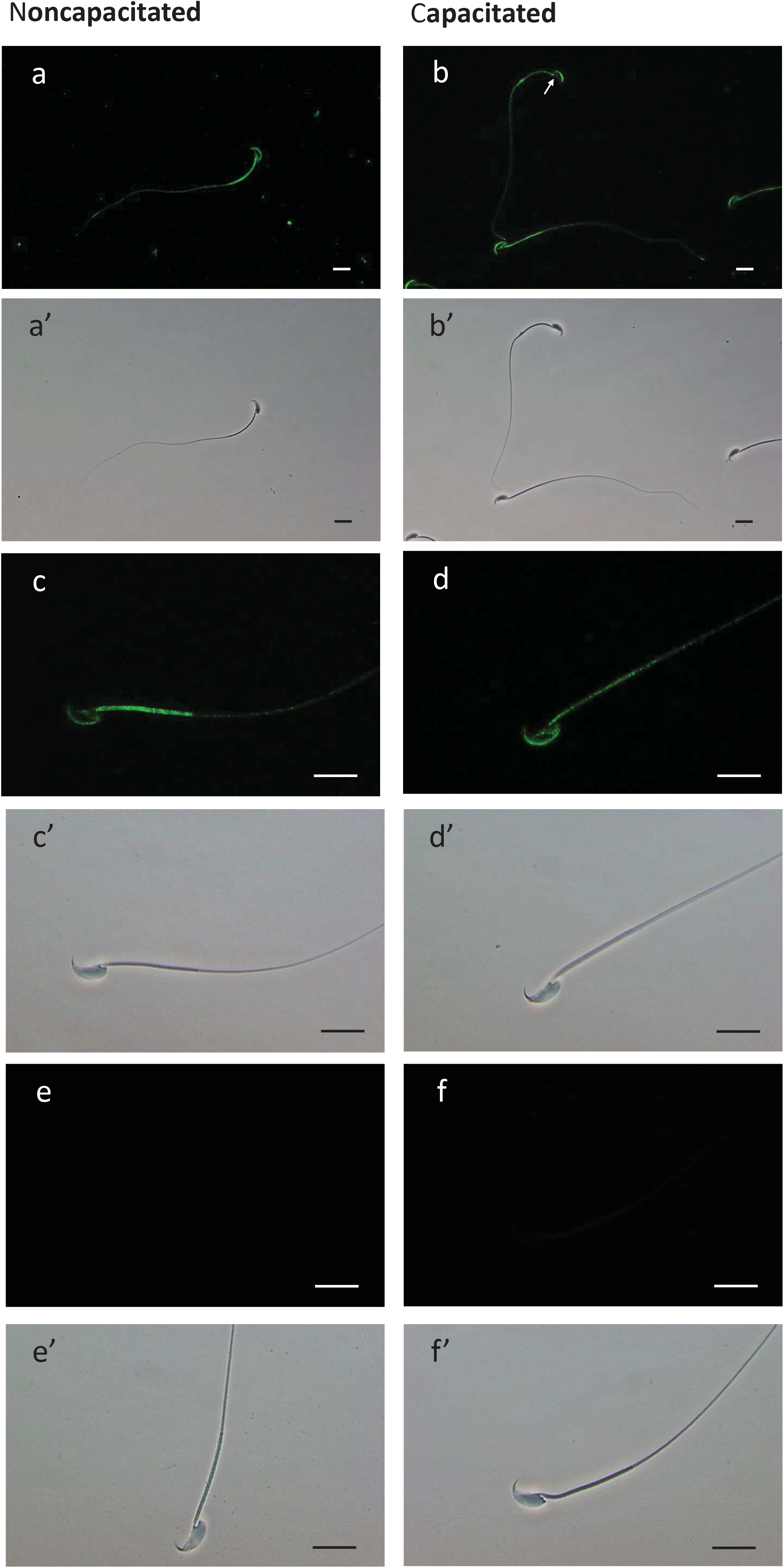
Immunofluorescent localization of EL and the corresponding phase-contrast micrographs of non-capacitated and capacitated mouse sperm cells. (**a-d’**) Fixed spermatozoa were immobilized on pre-coated slides, permeabilized with ice cold methanol and incubated overnight with anti-EL antibody. Alexa Fluor 488-conjugated secondary antibody was used for detection of the primary antibody. Primary antibody was omitted in control (e and f). The scale bars represent 20 μm. The experiment was repeated using four biological replicates.

### Loss in EL activity occurs during capacitation, independent of H89 and the acrosome reactions

In order to determine if the loss of Sia residue on EL affected enzyme activity, we specifically measured PLA_1_ activity on both non- and capacitated spermatozoa. Furthermore, as the immunofluorescence showed quite a varied pattern of EL expression, including acrosome location, we wanted to see if the loss of the acrosome had any effect of the overall EL activity. Therefore, we incubated sperm under non- or capacitating condition, with and without H89. Secondly, both non- and capacitated spermatozoa were induced to undergo the acrosome reaction. These sperm cells were subsequently washed, then the level of intact, partial or complete acrosome loss were measured using FITC-PNA staining. As shown in Figure 5, capacitated sperm plus ionophore A23187 had the highest level of complete acrosomal loss as expected.

**Figure 5.**
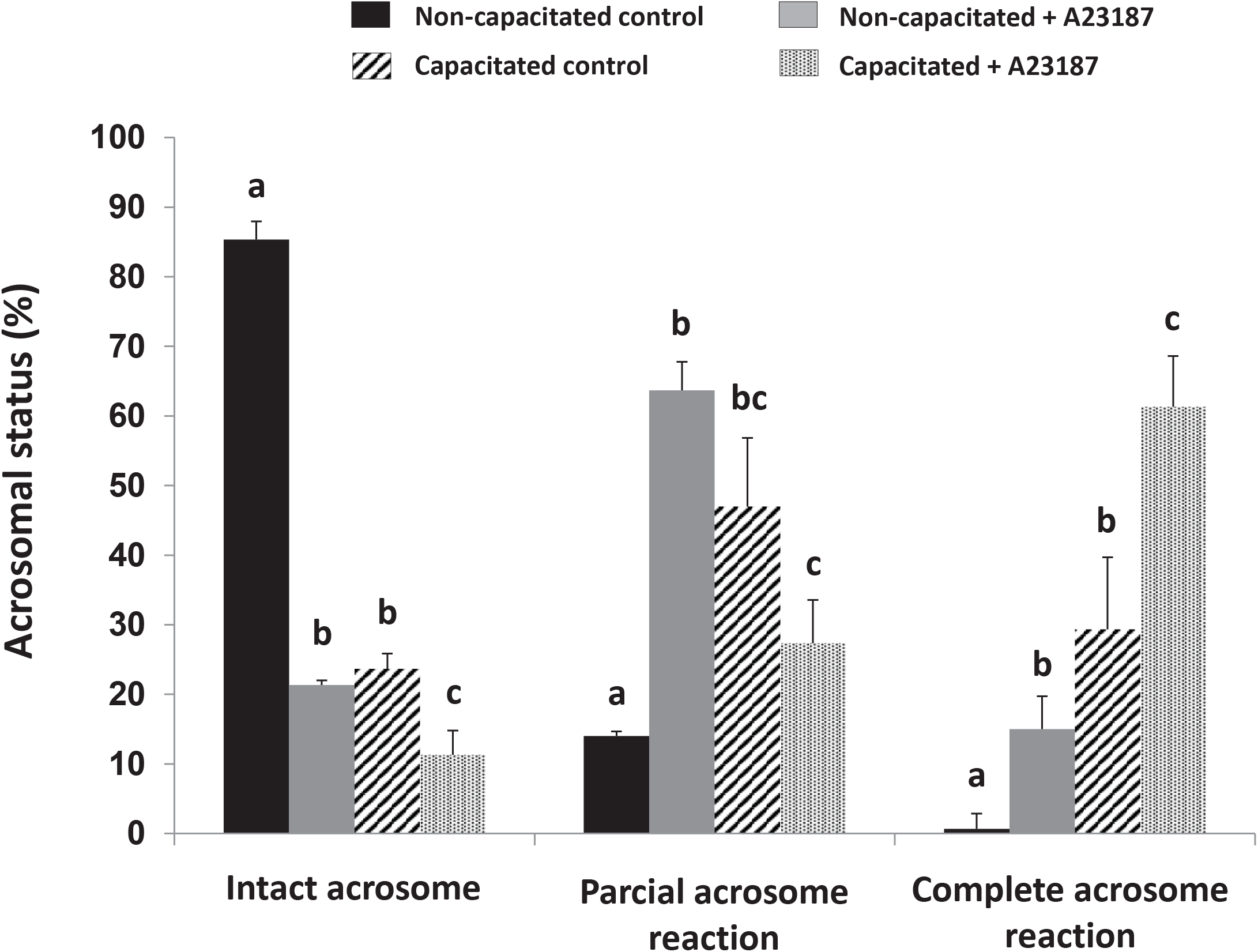
Assessment of acrosomal status by FITC-PNA labelling of non-capacitated (black and grey bars) and capacitated (hatched and dotted bars) mouse sperm either with (grey and dotted bars) or without (black and hatched bars) previous incubation with calcium ionophore A23187. Sperm were counted for intact, partial or complete acrosome reaction. The data were expressed as mean ± SE. Different letters represent statistical differences (p < 0.01) between treatments within each category (intact, partial or complete acrosome reaction). The graph represents the average of 5 biological replicates.

To determine whether a change in EL activity occurred during capacitation, we measured PLA_1_ activity before and after capacitation, together with the inhibitor H89 or the acrosomal-inducer, ionophore A23187. We observed a significant loss in the PLA_1_ activity of EL during capacitation (Fig. 6A, bars N-cap *vs* Cap). An example of the loss in PLA_1_ activity is demonstrated in Figure 6B. Here, two of the five biological replicates are shown over time from either non-capacitated (solid line) or capacitated (dotted line) sperm populations. Addition of the PKA inhibitor H89, which is commonly used to prevent capacitation, failed to abrogate the loss of PLA_1_ activity (Fig. 6A, N-cap + H89 *vs* Cap + H89). Furthermore, the loss of the acrosome had no further bearing on the reduction of the PLA_1_ activity in EL, with acrosome-reacted capacitated spermatozoa (Fig. 6A, bar Cap + A23187) having a similar activity to the capacitated (Fig. 6A, bar Cap) and the H89 “capacitated” (Fig. 6A, Cap + H89) cells. In all cases, the amount of activity was normalized to the amount of endothelial lipase present with an immunblot.

**Figure 6.**
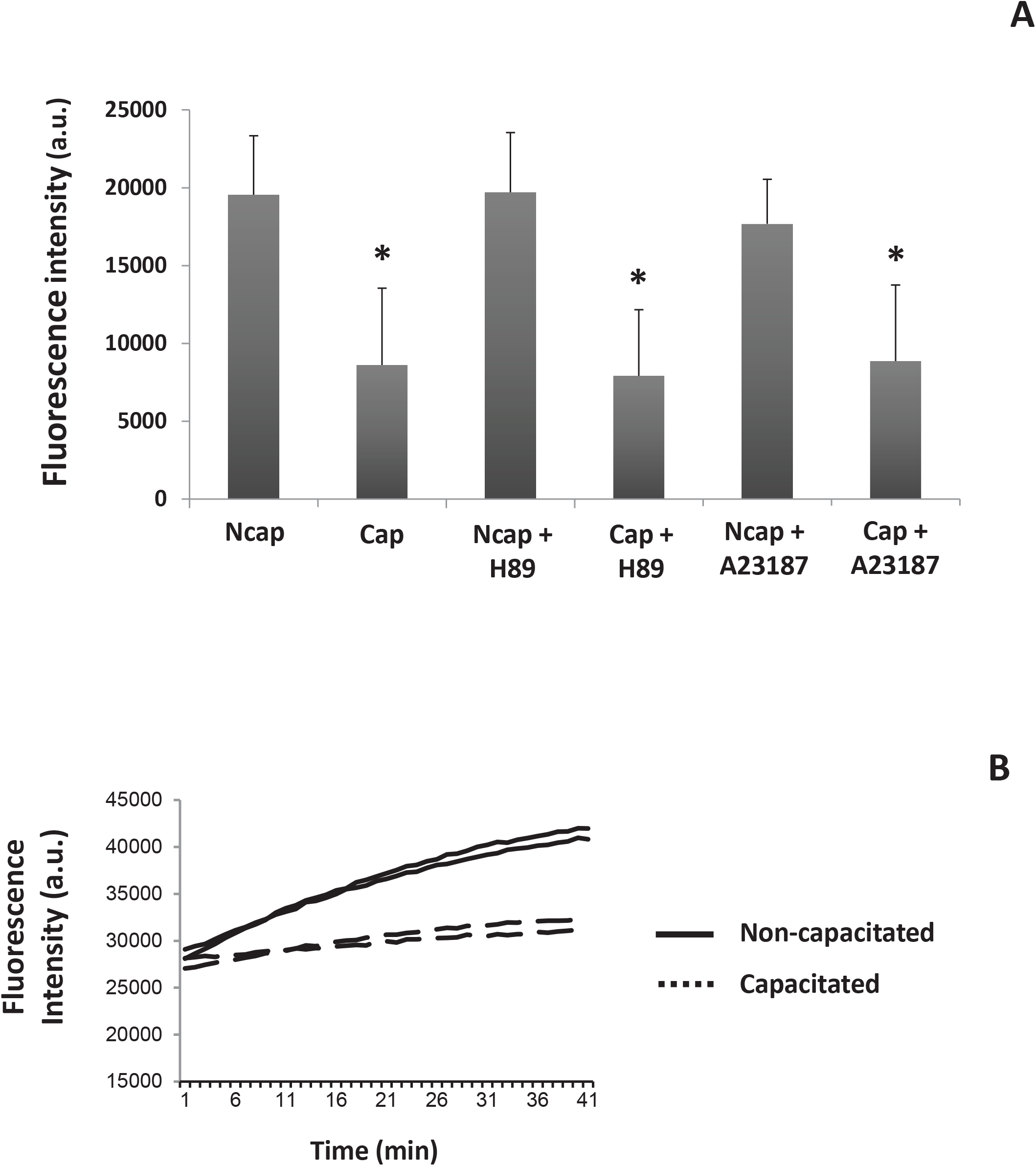
Phospholipase A_1_ (PLA_1_) activity and the effect of the addition of A23187 and the PKA inhibitor H89 on non- and capacitated mouse spermatozoa. Enzyme activity was measured using the fluorescent PED-A_1_ substrate. **(A)** PLA_1_ activity rate, within the first 10 min, for sperm cells previously incubated in non-capacitating or capacitating media with or without H89 and A23187. Image represents the mean value of five biological experiments. **(B)** Example of the raw data for two non-capacitated (solid line) and two capacitated (dotted line) samples.

### Sialylation of N612 inhibits ACO2 activity

Spermatozoa are catabolic in nature and, as such, it is not a surprise that the majority of Sia residues are lost during capacitation. However, in this study, we observed one enzyme that had an increase in sialylation, namely ACO2. To further understand this finding, we measured total Aconitase activity before and following mouse capacitation. As shown, during capacitation a statistically significant decrease in the level of Aconitase activity was observed (Fig. 7). Although this measurement would include both cytoplasmic and mitochondrial Aconitase activity, we reasoned that sperm have very little cytoplasm, therefore, the bulk of Aconitase activity should be from the mitochondrial form.

**Figure 7.**
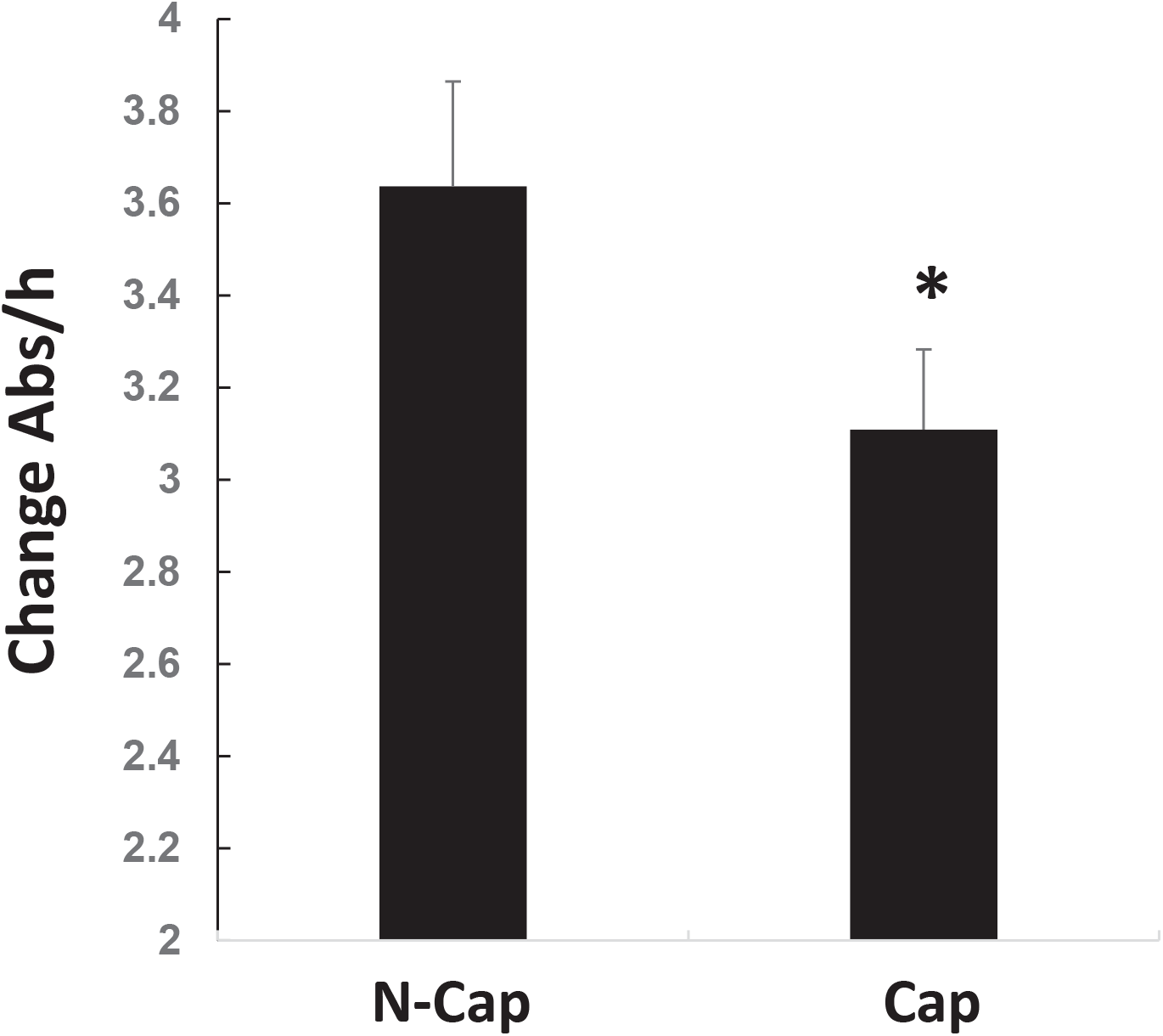
Decrease in Aconitase activity during capacitation. Spermatozoa from mice were retrieved and incubated under non- or capacitating conditions. Following protein extraction, Aconitase activity was measured. To ensure equal amounts of Aconitase were present, the lysate was precipitated, run into SDS page, transferred and probed using anti-Aconitase antibody. The data shown is the average (± SD) of four biological replicates. Asterisk represents statistical significance (p < 0.01).

To confirm that a decrease in ACO2 activity occurs specifically through sialylation at N612, we made both WT and Sia mimic, whereby the N612 was replaced by the negatively charged Aspartic acid (N612D). In both cases (WT and mimic), we made a GFP- and a HIS-tagged separate proteins.

Expression of the GFP-tagged proteins showed both WT and mutant were expressed in the mitochondria as expected (Fig. 8A-D). Of interest, under fluorescence microscopy, we also noted a decrease in the level of GFP-tagged mutant, yet the number of cells transfected was equal (compare Fig. 8A and 8B). To support this, we ran the cells through flow-cytometry. Analysis demonstrated that, in every replicate, the same number of cells were transfected (on average 35%), however, the mean level of GFP-fluorescence was significantly lower in the N612E-Aconitase expressing cells. For example, the histogram on Figures 8C and 8D demonstrates that the peak height for GFP fluorescence in WT is slightly higher (Fig. 8E, black downward arrow) than in the N612E mutant (Fig. 8F, black downward arrow). This suggests that cells are less favorable to the expression of the N612E mutant over the WT. To confirm this hypothesis, we ran a SDS gel, transferred and probed it with antibodies against GFP-(not shown) or (HIS)_6_-tagged proteins (Fig. 9A,B). As shown, WT ACO2 is more abundant than the mutant form (Fig. 9A,B; compare lanes 1 to 4). From our estimation, the N612E expression is between 25-50% of that of WT (see lanes 3 and 4 *vs* 5).

**Figure 8.**
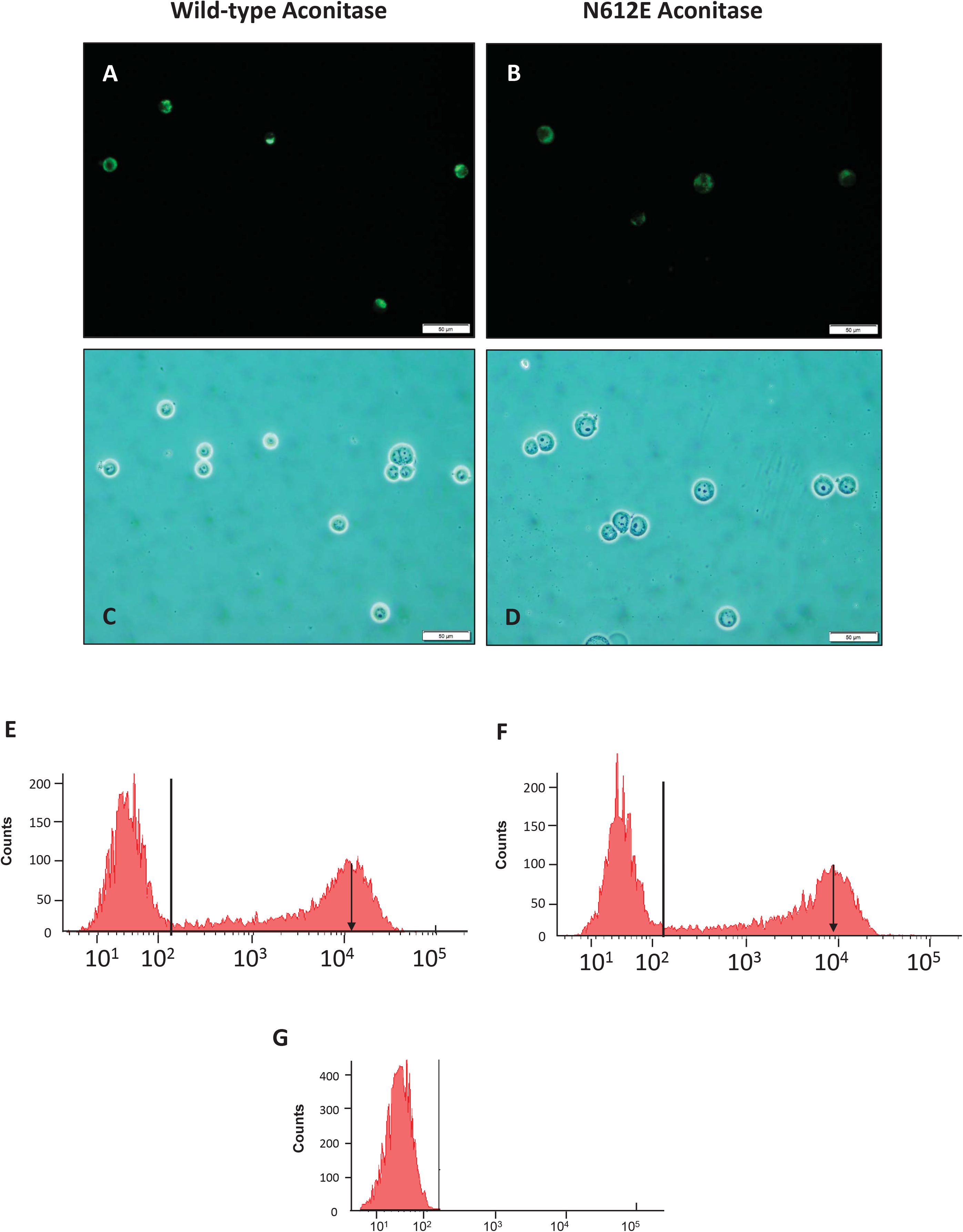
Following cloning and insertion of the Aconitase gene into plasmid, site directed mutagenesis was used to create the N612E mutant. **(A-D)** HEK293 cells were transiently transfected with GFP-tagged WT **(A,C)** or N612E mutant **(B,D).** Both anti-GFP tag fluorescent **(A,B)** and phase **(C,D)** images were taken. Scale bar = 50 μM. **(E-F)** Transfected cells were run through flow cytometer. Shown is the FL-1 channel vs counts of **(E)** GFP-tagged WT or **(F)** GFP-tagged N612E mutant. Arrow points to the mean signal intensity for each sample. Horizontal line represents the gate used for cell counting. **(G)** shows signal from non-transfected cells.

**Figure 9.**
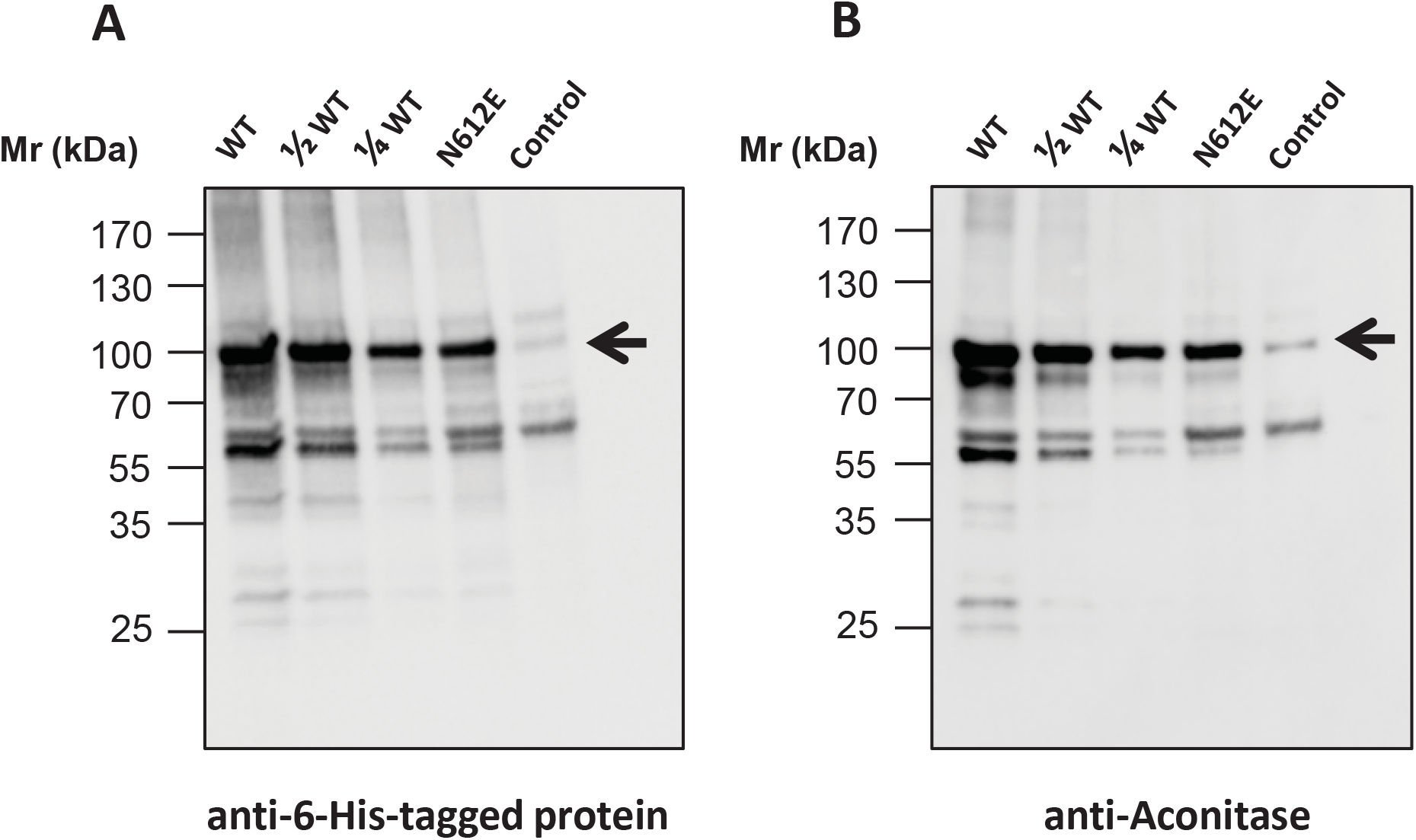
HEK293 cells were transfected with either (HIS)_6_-tagged WT or (HIS)_6_-tagged N612E mutant for Aconitase. Cells were lysed and recombinant Aconitase purified on nickel beads. Following elution, samples were run and probed with **(G)** anti-(HIS)_6_-tagged protein or **(H)** anti-Aconitase. Control samples comprise non-transfected cells.

We next measured the ACO2 activity of the WT and N612E mutant (Fig. 10). The use of equal amounts of recombinant protein was confirmed by immunoblotting. Remarkably, even when left overnight, we were unsuccessful in obtaining any ACO2 activity from the N612E mutant. In contrast, we could easily detect WT ACO2 activity. When put together, these datasets suggest that a switch in the ACO2 activity occurs during capacitation. This change in activity pattern is likely related to the modifications in the metabolic pathways described during the capacitation of mouse spermatozoa (i.e., from the oxidative phosphorylation pathway to the glycolytic pathway).

**Figure 10.**
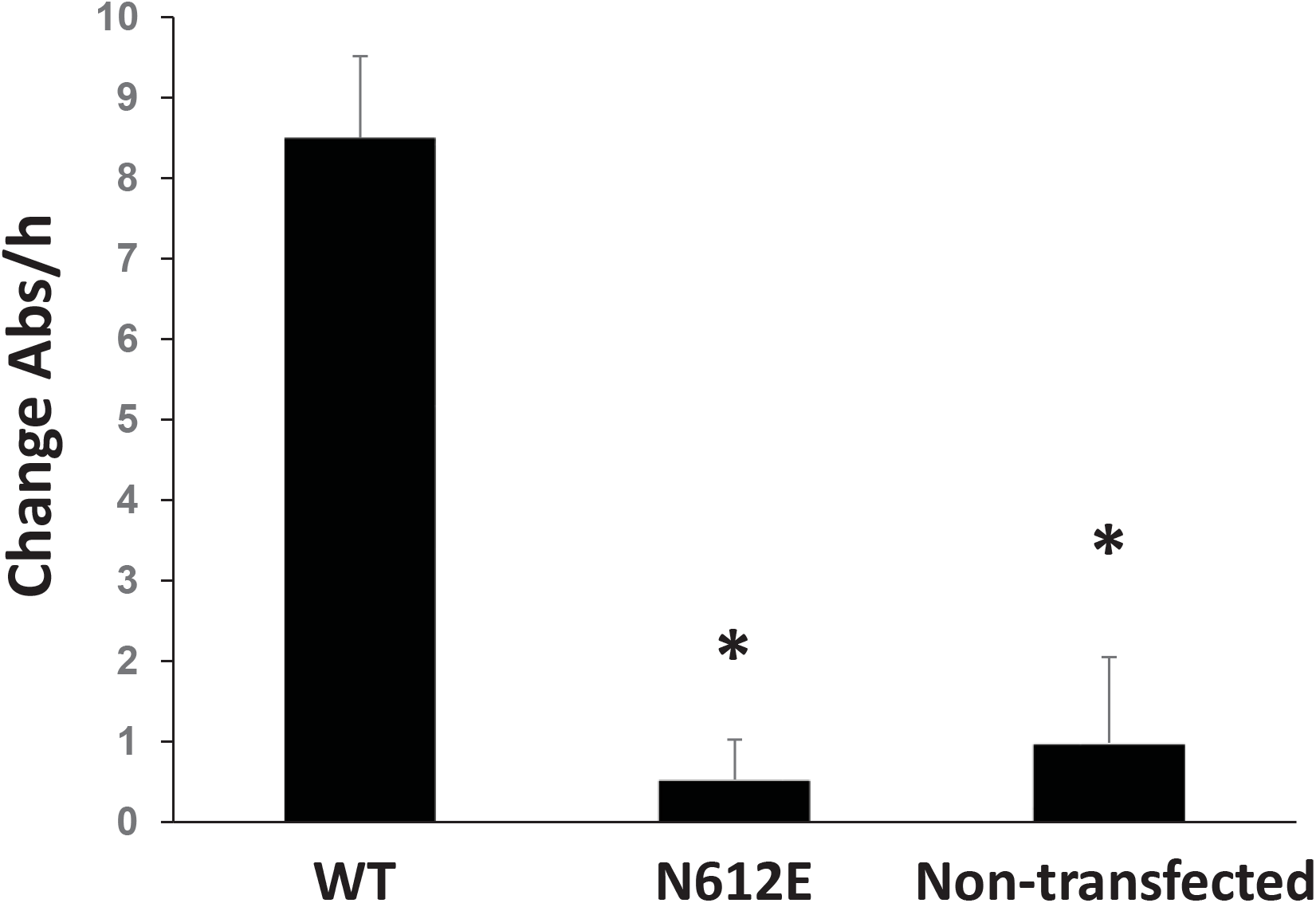
N612E-mutant Aconitase displays no activity. HEK293 cells were transiently transfected with (HIS)_6_-tagged WT and (HIS)_6_-tagged N612E mutant. Aconitase was purified over nickel beads, run in SDS-PAGE and the immunoblotting performed to ensure that equal amounts of protein were used. Bars demonstrate the average (n=6) activity of Aconitase with SD. Non-transfected cells were used as controls.

### Modelling the bound sialoglycoprotein to Aconitate hydratase

To understand the impact sialation would have on N612, we modelled the effect. Published crystal structures of Aconitate hydratase [45–48] and our highly homologous model, describes a large macromolecular structure comprising of four distinct domains employing a [4Fe-4S] cluster to catalyse the stereospecific dehydration of citrate to isocitrate [49]. Within the enzyme itself, sits several highly conserved amino acids, such as Asp192, His194 and Arg607 that are crucial for ACO2 activity (shown in Fig. 11). N612 resides on a short α-helical element immediately atop the active site, in an area often denoted as domain 4 [50]. In order to fit a sialic acid residue with N612, multiple changes in rotamers of sidechains were required. This allowed the formation of new salt bridges and hydrogen bonds between the sugar and the α-helical structure including the residue N614 (Asn615) and Gln563 which is present on an adjacent loop (Fig 11A,B). This data suggest that inhibition of Aconitase activity, through siltation of Aspargine 612, is due to major distortion of the active site which prevents catalysis from occurring.

**Figure 11.**
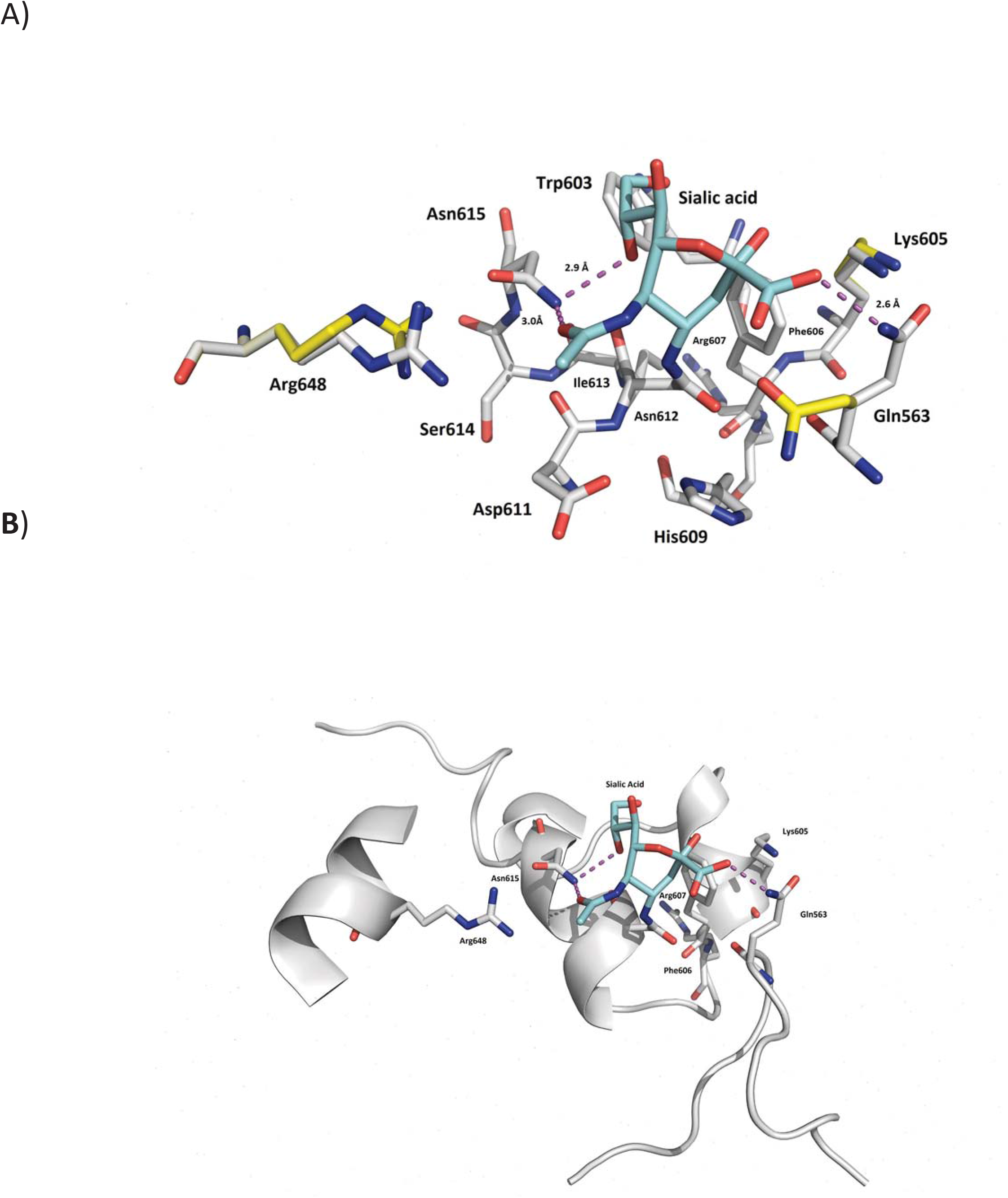
(A) Ball and stick together with (B) ribbon model of aconitase. The different amino acids are labelled as shown. The purple dotted line represents hydrogen bonding. Shown is sialic acid docked onto aspargine 612.

## Discussion

Despite the importance of capacitation, the molecular mechanisms underlying this process are not yet fully understood. Previous studies have suggested that one facet of capacitation is a loss in Sia residues, which may be modulated by one (humans) or two (mouse) neuraminidases, namely neuraminidase 1 and 3 [27]. In the present study, using a LC-MS/MS-based approach, we were able to investigate capacitation-related changes of N-linked glycoproteins bearing terminal Sia. Surprisingly, we found very little regulation of Sia within proteins groups following *in vitro* capacitation, with only 6.3% (9 of 142 peptides) demonstrating a significant change.

According to our data, the enzyme EL is one of the sperm proteins in which the Sia content is altered during capacitation. Previous studies have shown that four (N62, N118, N375, and N473) of the five potential N-glycosylation sites of human EL are occupied by glycan moieties [51, 52]. In the present study, the sugar moiety at the N62 glycosylation site of EL was found to contain Sia in its structure. In addition, the peptide containing this glycosylation site was significantly reduced after *in vitro* capacitation, despite the fact that both quantity and molecular weight of EL remained unchanged. This suggests that a loss of a small glycan moiety or of a Sia residue itself occurs within EL during capacitation. Notably, a sialylated N-glycan structure at the corresponding glycosylation site (N64) has also been identified by our group in the EL of rat sperm [33]. In this case, spermatozoa were taken from the caput, corpus and cauda regions of the epididymis. Remarkably, the Sia residue within EL was only found in spermatozoa derived from the caudal location. Furthermore, we have observed that the amount of EL within rat spermatozoa does not change, suggesting that Sia is added to EL during epididymal transit (**data not published;**[33]). Given that Sia residues are removed from the same glycosylation site during capacitation, it is likely that this glycan moiety plays a specific role in regulating EL enzyme activity.

The glycosylation site at N62 of mouse EL is a conserved feature among animals and other members of the triglyceride lipase gene family, such as lipoprotein lipase and hepatic lipase. Using recombinant proteins, two separate studies have produced point mutations of the amino acids that are glycosylated in EL. Interestingly, in both cases, the loss of N62 led to increased EL activity [51, 52]. Due to their negative charge and hydrophilicity, Sia residues within this glycan moiety could influence the structure and/or substrate specificity of EL, therefore, regulating its enzymatic activity. Of note however, we observed a decrease in the PLA_1_ activity of EL following reduction of its sialylated glycopeptides (N62 glycosylation site) during capacitation. We can only assume that, besides the loss of N62, EL is likely to be regulated in other (as yet unknown) ways in order to switch off its activity.

In addition to EL, we found N612 sialytion of ACO2 increase following *in vitro* capacitation. This enzyme catalyzes the non-redox reaction of the TCA cycle in which stereo-specific isomerization of citrate to isocitrate occurs [53]. Adequate supply of ATP is essential to support capacitation-associated changes such as hyperactivation [54]. In mouse, it is fairly well understood that, during capacitation, there is a switch from oxidative phosphorylation over to glycolysis [5]. Thus, non-capacitated sperm show high oxygen consumption, which diminishes as sperm make the switch over to glycolysis during capacitation [5]. Herein, such a switch could be brought about through sialylation of ACO2 particularly N612. Indeed, modelling of the enzyme suggests that the Asn group sits atop of the ACO2 activity site (Fig. 11) in a highly conserved region. Analyses of multiple X-ray crystal structures of Aconitate hydratase has shown that a vast array of residues from all four domains of the enzyme are required to carry out catalysis [55] whether it’s to bind and recognize substrate or ligate the [4Fe-4S] cluster. Similarly, this complex array, dependent on bound ligands, displays a number of nuanced and obvious conformational changes within its active site and other domains [55]. Thus, it is apparent that in our model, inhibition likely occurs through segmental conformational change of the antiparallel helical motif that N612 is a part (residues 606-612 in our model; Fig. 11). As this includes active site residue Arg607, it is possible changes in substrate binding may occur to prevent catalysis such as a rotameric shifts or larger domain movements that may prevent substrate binding and release, a trait often linked with N-linked glycosylation [56]. With the observation that both the mutation to Glutamate (Glu612) and the Asn612 sialic acid (N-link) modified enzymes shuts down activity of the enzyme and the former modification in our models (not depicted) suggests a new salt bridge or hydrogen bond with the either the sidechain amine of Gln563 or Lys605 is likely. This further underlines the potential sensitivity of ACO2 activity with respect to changes to this motif with the potential they instigate larger downstream changes in other domains.

Inspection of Uniprot suggests that the glycosylation at N612 has never been reported in any other cell type and, as such, may represent a novel mechanism, attributed just to sperm cells. Unfortunately, compounds to block or inhibit sialic acid transferases are not cell permeable and, for this reason, we were unable to directly ascribe the significance of a lack of ACO2 activity to sperm physiology. Additionally, an indiscriminate blockage of sialyltransferases would have raised doubts about what other proteins/pathways could also have been co-inhibited and this obfuscated the interpretation. Our leading hypothesis is that glycolysis is required for the rapid movement of the sperm flagella, in a process known as hyperactivation. The latter being essential for fertilization to occur. Therefore, sperm cells being transcriptionally and translationally silent cells may change ACO2 activity via sialylation of this protein, which facilitates the shuttling of its metabolic process from oxidative phosphorylation over to glycolysis.

## Abbreviations

ACN: acetonitrile
Asn: asparagine
dbcAMP: dibutyryl-cAMP
EL: endothelial lipase
LC-MS/MS: liquid chromatography tandem-mass spectrometry
MS: mass spectrometry
MAM: meprin/A5 antigen/mu receptor tyrosine phosphatase
ACO2: mitochondrial aconitate hydratase
PNGase F: peptide-N-glycosidase F
PLA_1_: phospholipase A_1_
PED-A_1_: phospholipase A_1_ selective substrate
Ser: serine
Sia: sialic acid
T: threonine
D: Aspartic acid
TiO_2_: titanium dioxide
TCA: tricarboxylic acid
TFA: trifluoroacetic acid
WT: wild-type ACO2 transfected

## Notes

This work was supported by the Brazilian National Council for Scientific and Technological Development (CNPq).

